# Saturation variant interpretation using CRISPR prime editing

**DOI:** 10.1101/2021.05.11.443710

**Authors:** Steven Erwood, Teija M.I. Bily, Jason Lequyer, Joyce Yan, Nitya Gulati, Reid A. Brewer, Liangchi Zhou, Laurence Pelletier, Evgueni A. Ivakine, Ronald D. Cohn

## Abstract

Over the last decade, next generation sequencing has become widely implemented in clinical practice. However, as genetic variants of uncertain significance (VUS) are frequently identified, the need for scaled functional interpretation of such variants has become increasingly apparent. One method to address this is saturation genome editing (SGE), which allows for scaled multiplexed functional assessment of single nucleotide variants. The current applications of SGE, however, rely on homology-directed repair (HDR) to introduce variants of interest, which is limited by low editing efficiencies and low product purity. Here, we have adapted CRISPR prime editing for SGE and demonstrated its utility in understanding the functional significance of variants in the *NPC1* gene underlying the lysosomal storage disorder Niemann-Pick disease type C1 (NPC). Additionally, we have designed a genome editing strategy that allows for the haploidization of gene loci, which permits isolated variant interpretation in virtually any cell type. By combining saturation prime editing (SPE) with a clinically relevant assay, we have functionally scored and interpreted 256 variants in *NPC1* haploidized HEK293T cells. To further demonstrate the applicability of this strategy, we used SPE and cell model haploidization to functionally score 465 variants in the *BRCA2* gene. We anticipate that our work will be translatable to any gene with an appropriate cellular assay, allowing for more rapid and accurate diagnosis and improved genetic counselling and ultimately precise patient care.

## Introduction

Precision or individualized medicine is necessarily predicated on a robust understanding of the genetic variation found in the population. Accordingly, the preponderance of VUS found in human disease genes represents a significant hurdle to the realization of precision medicine. Traditional methods for resolving the functional or clinical significance of a VUS, such as co-segregation studies or post-hoc functional assays, are limited in scale. As both whole-exome and whole-genome sequencing become more common, particularly in clinical settings, the number of VUS is rapidly accumulating which necessitates accurate scalable methods of variant interpretation^1^. To this end, there has been remarkable technology development in multiplexed assays of variant effects (MAVEs). These assays allow for the simultaneous interrogation of functional effect of all possible nucleotide or amino acid substitutions using diverse phenotypic readouts^2,3^. Importantly, these assays are proactive, often functionally classifying mutations before they are seen in the clinic. These assays, however, often uncouple the variants-of-interest from their native genomic context being overexpressed as cDNAs from artificial cassettes or in non-human cell types^4,5^. This has been addressed by saturation genome editing (SGE), which uses CRISPR-Cas9 genome editing to introduce all possible SNVs into a targeted region of the endogenous genomic locus^6–8^.

To date, SGE has required HDR to introduce the saturating sets of genome edits. Typically, however, HDR is inefficient and results in an excess of indel formation due to competing non-homologous end-joining. There have been demonstrations of near-saturation genome editing using CRISPR base editing technologies, though these are inherently limited to transition point mutations and the targetable area of a gene is restricted by protospacer adjacent motif (PAM) location and availability^9,10^. Furthermore, the introduction of unintended on-target bystander mutations in the wide editing window can complicate variant interpretation in base editing experiments^10^. The recently developed CRISPR prime editing system represents an attractive addition to the SGE toolkit. Prime editing allows for the precise introduction of all possible nucleotide substitutions alongside small insertions or deletions^11^. In contrast to HDR, prime editing does not require exogenous donor templates and provides predictable editing outcomes and high product purity. In addition, though SGE has been recently expanded to the diploid TMD8 cell line^8^, previous studies have relied on the near haploid HAP1 cell line^6,7,12^. While HAP1 cells simplify interpretation by allowing for functional analysis of variants in isolation, they limit SGE to genes expressed or phenotypes elicitable in these cells.

Here, we sought to adapt the recently developed CRISPR prime editing system to effectuate SGE. Furthermore, we developed an allele-specific genome editing strategy that leverages naturally occurring genetic variation to generate ‘haploidized’ gene loci. We posit that these two modifications will generalize SGE ultimately widening its adoption and usefulness. We primarily focused on the NPC intracellular cholesterol transporter 1 gene (NPC1), which underlies the rare lysosomal storage disorder Niemann-Pick disease type C1^13^. NPC1 is required for proper cholesterol trafficking and pathogenic mutations in *NPC1* lead to an accumulation of cholesterol in late endosomes and lysosomes, ultimately resulting in cell death. While NPC is clinically heterogenous^14^, the disorder is characterized by progressive neurodegeneration that leads to premature fatality^13^. Patients with NPC and their families typically face protracted diagnostic odysseys, due in part to a poor understanding of the clinical significance of most variants in *NPC1*^13,14^. These VUS delay treatment administration and complicate genetic counselling efforts in NPC. Accordingly, a proactive mapping of the functional consequences of variants in *NPC1* address an unmet clinical need. We developed and validated a novel fluorescence-activated cell sorting (FACS) pooled functional assay for variants in *NPC1*, demonstrating the first instance of SGE not dependent on cell-growth or essentiality phenotypes. We combine this functional assay with SPE in a *NPC1* haploidized HEK293T model, functionally scoring and classifying 256 unique variants. We show that these scores are strongly correlated with *in silico* predictions of pathogenicity and that they reflect specific molecular mechanisms underlying the pathogenesis of each variant. To illustrate the versatility of haploidized cell models and SPE, we extended our analysis to the clinically actionable tumor suppressor gene, *BRCA2*. Together, we functionally assay 721 variants across these two relevant disease genes in haploidized HEK293T cells. We show that SPE is characterized by high and uniform editing efficiencies and relatively low indel formation. We expect that SPE and haploidized cell models will be of broad utility to the functional genomics community and we anticipate the functional classifications made in the *NPC1* and *BRCA2* genes will aid in establishing a clinical diagnosis that will improve genetic counselling and carry the potential toward a more individualized management and treatment approach of affected individuals.

## Results

### Allele-specific genome editing allows for haploidization of gene loci

In a diploid cell line, variant effect can be masked by heterogeneous editing events, posing a challenge for accurate functional interpretation. To circumvent this, initial demonstrations of saturation or near-saturation genome editing relied on the HAP1 cell model, a near haploid human cell line^6,7,12^. The HAP1 cell line, however, can spontaneously revert to a diploid state in culture and has required secondary genetic manipulations to yield sufficient editing efficiencies^7^. Furthermore, continued reliance on HAP1 cells fundamentally limits the applicability of SGE to genes expressed or phenotypes elicitable in this cell line. To address this, we sought to generate a non-HAP1 model system that allows for isolated variant effect analysis. We hypothesized that “haploidized” gene loci could be generated by deleting genomic regions encompassing a large fragment of a gene-of-interest in an allele-specific manner. Specifically, we sought to design a pair of guide RNAs, each specific to a naturally occurring heterozygous single nucleotide polymorphism (SNP) that span the *NPC1* gene, that when delivered simultaneously, would result in a deletion of the intervening segment. We reasoned that if sufficient specificity towards these polymorphisms could be achieved, editing would occur only on the targeted allele, leaving the non-targeted allele unmodified.

Using existing whole-genome sequencing data from the HEK293T cell line^15^, we selected one SNP found in exon 6 –*NPC1* c.644A>G – and another SNP found in the intron following exon 24 –*NPC1* c.3754+34A>G – for haploidization (Figure 1A). A relative copy number of 2:1 of G:A at each site was estimated by Sanger sequencing, indicating that the parental HEK293T cell line was triploid at the *NPC1* locus. We expected that the third copy of NPC1 was from whole-or partial-chromosomal duplication and thus the ‘G’ sites were on the same chromosome. Accordingly, CRISPR-Cas9 targeting of these alleles would result in the deletion of the genomic segment intervening these SNPs. The exon 6 SNP occurred in the first PAM-proximal nucleotide of a potential *Streptococcus pyogenes* Cas9 (SpCas9) guide sequence. We designed a guide RNA that was complementary to the two ‘G’-containing targeted alleles and was mismatched to the non-targeted ‘A’-containing allele. The intronic SNP resulted in a non-canonical^16^ ‘NAG’ SpCas9 PAM on the non-targeted allele and a canonical ‘NGG’ SpCas9 PAM on the two targeted alleles. We hypothesized that the mismatch in the guide seed-region^17–19^ would prevent cleavage of the non-targeted allele at the exon 6 SNP and that the non-canonical PAM present on the non-targeted allele would diminish cleavage at the intronic SNP.

**Figure 1.**
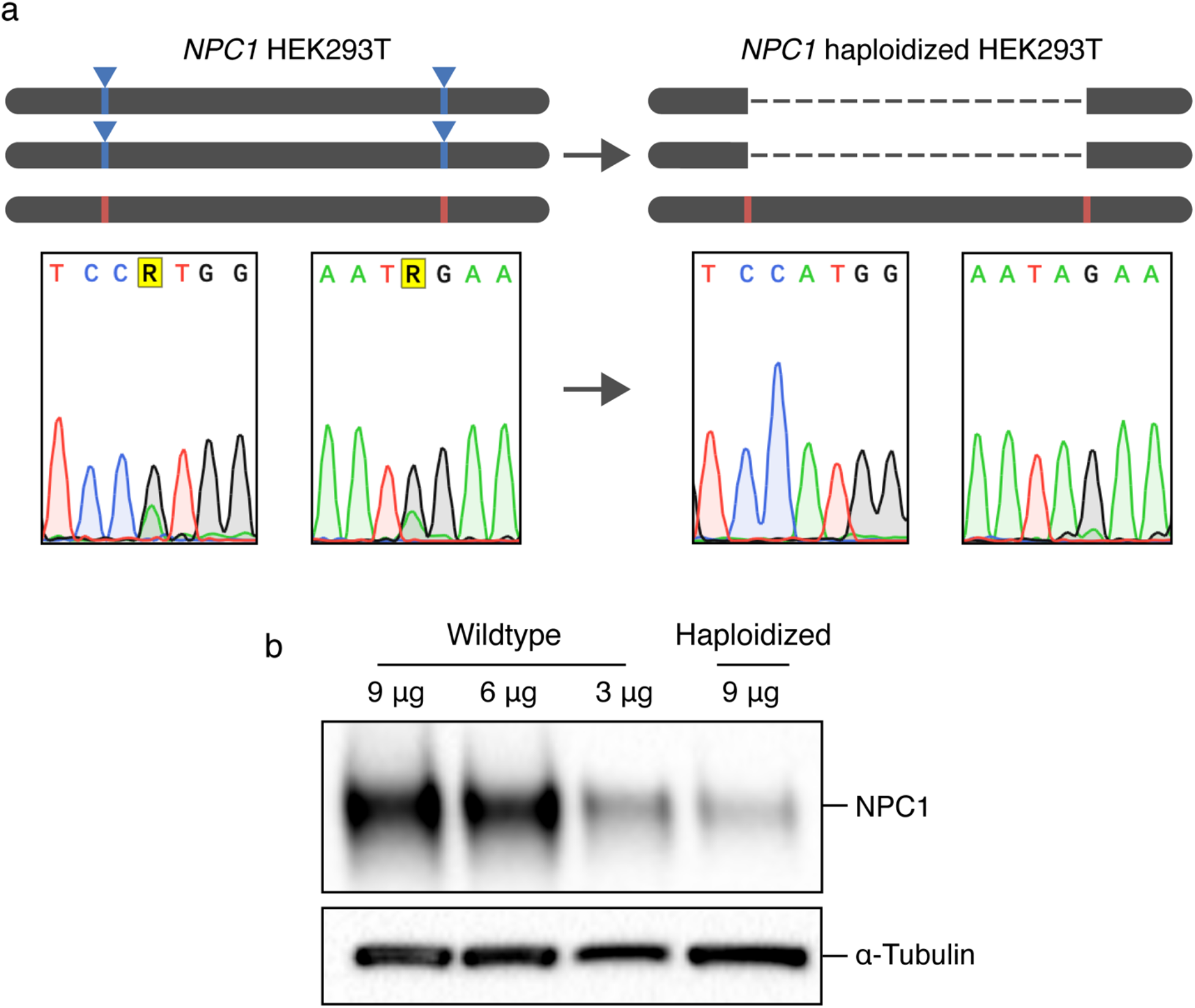
Allele-specific genome editing permits haploidization of the *NPC1* locus. (A) Top, schematic overview detailing the haploidization approach, bottom, sequencing reads from HEK293T cell lines pre- and post-haploidization. (B) Approximately 1/3^rd^ of NPC1 protein expression is detectable in *NPC1* haploidized HEK293T cells compared to wildtype controls.

We delivered these two guides on separate vectors both co-expressing SpCas9 and isolated clonal populations from single cells. We genotyped each clone by first attempting to amplify the deletion junction that was formed between the two cut sites, then amplifying and sequencing the two SNP target sites. In the initial attempt to derive the haploidized cell line using this strategy, all clones harbouring a deletion between the two guide RNA sites had small indels at the exon 6 target-site on the non-targeted allele, indicating that the single nucleotide mismatch between the guide RNA and the non-targeted allele was insufficient to protect against cleavage. The non-canonical PAM present on the non-targeted allele, however, did diminish cleavage activity as this target site remained unmodified in all derived clones. Recent work has demonstrated that unwanted cleavage at sequences similar to the desired target-site can be minimized by targeting the former with catalytically-inactivating truncated guide RNAs, or dead guide RNAs^20,21^. These dead guide RNAs allow for Cas9 binding but not cleavage, protecting the site from modification. We designed a 15-nucleotide dead guide RNA where the ‘A’ present on the non-targeted allele was in the 5^th^ PAM-proximal position. Finally, we codelivered plasmid expression vectors for the three described guide sequences, each coexpressing SpCas9 and subsequently isolated monoclonal cell lines. The addition of the dead guide RNA significantly reduced undesired indel formation on the non-targeted allele. In the five clones isolated with the desired deletion on the two targeted alleles, the non-targeted allele had no modification at either polymorphic cut site. We performed digital droplet PCR to determine copy number of the deleted region to ensure it was not retained elsewhere in the genome and found that three of the five clones had the deleted region in only a single copy. We performed filipin staining on these haploidized clones and confirmed that cholesterol homeostasis was maintained with *NPC1* in a single copy (Supplementary Figure S1). Haploidization of *NPC1* was reflected in protein expression, with edited clones showing approximately 1/3^rd^ the amount of total NPC1 expression compared to the parental HEK293T cell line (Figure 1B). While fragments of the *NPC1* gene remain on the two targeted alleles, it is unlikely that their expression functionally contributes to cholesterol trafficking. The segment removed from the two targeted alleles encodes three of the five protein domains of NPC1, including the functionally essential sterol sensing and NPC2 binding domains. Altogether, this haploidized model simplifies the interpretation of *NPC1* variants by allowing for functional and molecular interrogation in effective isolation. Furthermore, the specific approach applied to haploidize this locus is not specific to either *NPC1* or HEK293T cells. Given the ample variation found in any human cell line, this strategy represents the basis for a generalized approach for the isolation of gene loci for functional studies of variant effect.

### Establishing a functional assay for NPC1 variant interpretation

The accumulation of cholesterol that results from a deficiency in NPC1 leads to the expansion of the lysosomal and late endosomal compartments^22^. This expansion can be visualized with the cell-permeable fluorescent dye LysoTracker™^23–25^. We hypothesized that we could functionally distinguish variants by measuring relative LysoTracker™ accumulation in a fluorescence-activated cell sorting assay. First, we used prime editing in the *NPC1* haploidized cell line to generate clonal models of two well-characterized mutations associated with NPC – NPC1 p.I1061T and NPC1 p.P1007A – alongside an NPC1 knock-out model harbouring the nonsense mutation NPC1 p.C909X. We first confirmed that a qualitative difference was apparent between the NPC1 p.C909X knockout cell line and wildtype controls when stained with LysoTracker™ (Figure 2A). While little is understood about how specific mutations contribute to the varied clinical manifestations and disease progression found in NPC, NPC1 p.I1061T has been associated with the classical juvenile onset NPC subtype and more rapid disease progression^26^ while NPC1 p.P1007A is recognized as a biochemical variant typically associated with later disease onset^27,28^. Of note, diagnostic filipin staining of patient fibroblasts that are heterozygous or homozygous for the NPC1 p.P1007A can result in inconclusive clinical interpretations due to mild or indiscernible cholesterol accumulation^29,30^. Indeed, these clinical differences are reflected in protein expression in both patient-derived fibroblasts^27^ and in the cell models we generated (Figure 2B).

**Figure 2.**
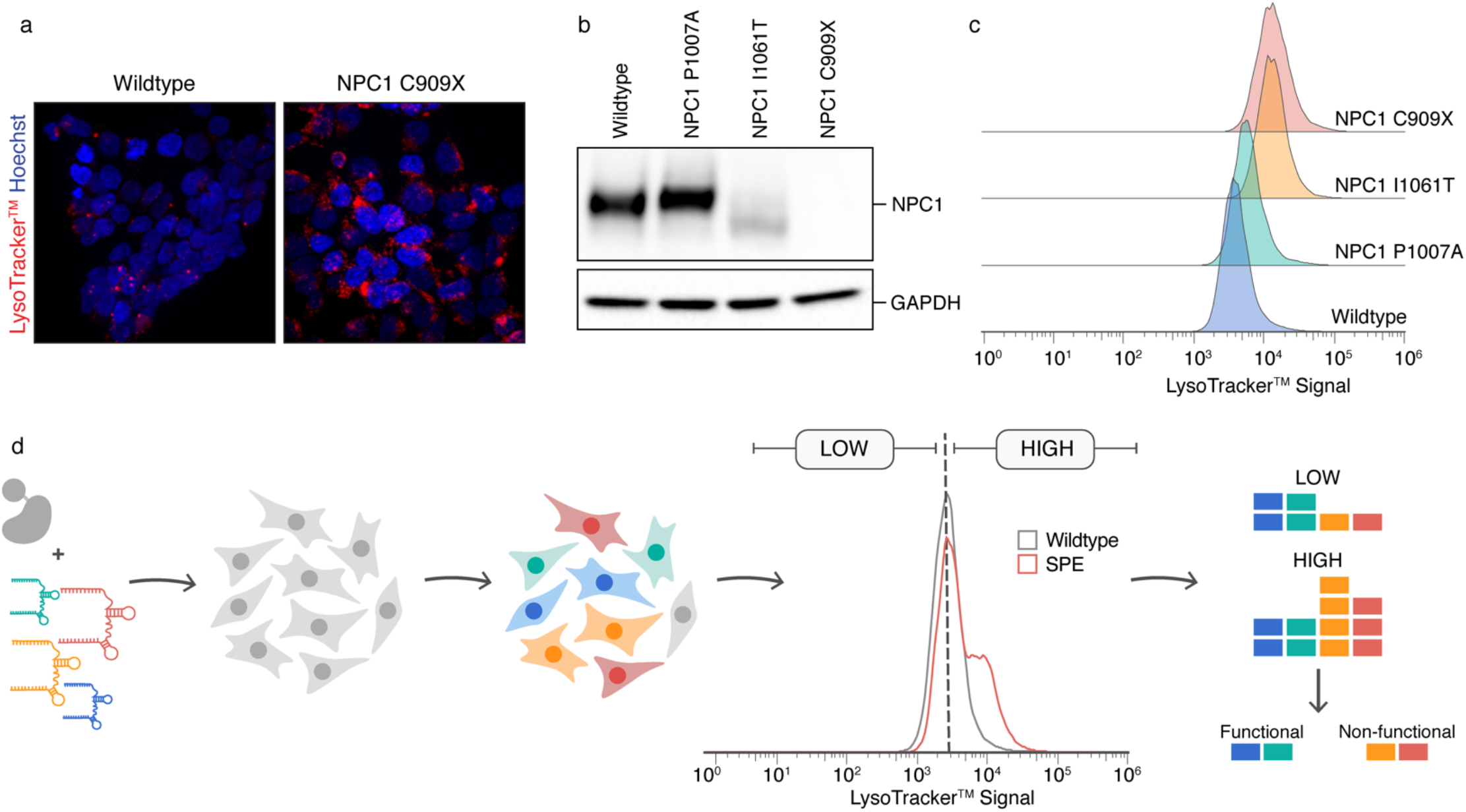
Establishment of a functional assay for variants in *NPC1*. (A) NPC1-deficient cells display increased LysoTracker™ accumulation compared to wildtype controls. (B) Immunoblot illustrating the varied protein expression patterns found in different *NPC1* variants in *NPC1* haploidized HEK293T cells. (C) Fluorescent distributions derived from fluorescence-activated cell sorting of LysoTracker™-stained mutant cell lines. (D) Schematic overview of the pooled functional assay for variants in *NPC1*.

To understand the sensitivity of a potential FACS-based functional assay for *NPC1* variants, we assayed the fluorescent distribution of each of mutant cell line compared to the haploidized parental HEK293T cell line when stained with LysoTracker™. Each mutant had a fluorescent distribution shifted right of the haploidized wild type, demonstrating that LysoTracker™ staining intensity can serve as a surrogate for the functional significance of variants in *NPC1* (Figure 2C). Encouragingly, the degree to which the distribution of LysoTracker™ signal was right-shifted corresponded to the existing clinical correlations described above, with NPC1 p.P1007A having a less pronounced shift relative to haploidized wild type compared to NPC1 p.I1061T and NPC1 p.C909X. These results support the use of a FACS-based assay for the purpose of functionally assessing variants in *NPC1*. Of note, this assay could be used to not only discriminate between pathogenic and benign variants but may also inform the degree of functional defect of a given pathogenic variant.

### Saturation prime editing of select regions of NPC1

Genomic edits made by prime editing are encoded in the reverse transcription (RT) template portion of the 3’-extension of the prime editing guide RNA (pegRNA)^11,31^. Accordingly, the region encompassed by the RT template can be mutated to saturation by delivering a library of pegRNAs, each encoding one of the possible nucleotide substitutions across the length of the editable region. To demonstrate the feasibility of saturation prime editing, we designed libraries of pegRNAs targeting four different regions across the coding region of *NPC1*. The pegRNA architecture differed for each site, with each having a primer binding site (PBS) 13 nucleotides in length and a RT template of either 13, 14, 22, or 43 nucleotides in length.

When initially designing and evaluating pegRNAs, we reasoned that the prime editing efficiency of a pegRNA might be correlated to the targeting efficiency of a normal guide RNA. This has since been confirmed by a recent study that defined the properties of pegRNAs correlated with efficient editing^32^. Accordingly, we chose six potential spacer sequences based on high computationally predicted editing efficiency. We constructed these pegRNA sequences with 3’-extensions composed of 13 nucleotide primer binding sites and 13 nucleotide RT templates. We evaluated each candidate 3’-extension by attempting to program a nucleotide substitution six base pairs downstream of the prime editor’s pegRNA nicking site, which would ultimately destroy the PAM sequence of that pegRNA. Four of the six pegRNA sequences produced editing efficiencies detectable via Sanger sequencing and were carried forward for further analysis. To maximize the editable window with these pegRNAs we arbitrarily chose and tested RT templates lengths ranging from 13 to 45 nucleotides. In line with previous research, we generally found that editing efficiencies greatly diminished with increasing RT lengths^11,32^. For the pegRNA architectures ultimately used, we chose the longest RT template length that supported estimated editing efficiencies greater than 20% as determined by Sanger sequencing of edited cell populations. To maximize editing efficiency, we co-delivered a nicking guide RNA during both the evaluation of candidate pegRNAs and in subsequent multiplexed SPE experiments. Each library was designed to include a silent PAM-destroying substitution alongside each targeted mutation, except when the targeted mutation itself disrupted the PAM sequence. By disrupting the PAM sequence, the initial successful edit prevented subsequent retargeting and thereby prevented multiple editing events occurring in the same cell. In addition, we included up to four pegRNAs in each library which encoded multinucleotide variants that resulted in nonsense mutations to serve as internal controls of our functional scoring. We generated these plasmids libraries and transfected them individually into the *NPC1* haploidized HEK293T cells alongside two expression vectors, one encoding the SpCas9 prime editor and another encoding a nicking guide RNA and a puromycin selection cassette. Each SPE experiment was carried out with 8 experimental replicates. After transfection, we used antibiotic selection for 72 hours to enrich for successfully transfected cells. To maximize potential editing efficiencies, we performed our FACS-based functional assay on each experimental replicate 10-days post-transfection. Based on our observation that different pathogenic mutants have varying degrees of excess LysoTracker™ accumulation, we devised a gating strategy based on the fluorescent distribution of an unedited control population. Specifically, our low fluorescence gate encompassed the bottom ~50% of the unedited control fluorescence distribution and the high fluorescence gate encompassed the top ~50% of the unedited control fluorescence distribution. Accordingly, we would expect functional variants to be equally represented across the two gates, while non-functional variants would be disproportionately represented in the high fluorescence gate (Figure 2D). We extracted the total genomic DNA from each gated population and sequenced the targeted area. We derived a function score for each mutation modelled which was based on the log-fold change from the low fluorescence gate to the high fluorescence gate. We pooled these scores from the four different libraries and scaled them such that the average nonsense score was approximately equal to 0 and the unmodified wildtype allele score was approximately equal to 1. The normalized function scores were highly reproducible across the 8 experimental replicates (Pearson’s *r* between 0.94 and 0.99, Supplementary Figure S2A). Next, we performed unsupervised clustering (Methods) to define the functional cluster and classified everything with a function score lower than that cluster as nonfunctional.

Altogether, the function scores from the four libraries encompassed *NPC1* c.1040-1052, *NPC1* c.1973-1983, *NPC1* c.2720-2758 and *NPC1* c.3091-3109 (Figure 3A, Supplementary Table 1). We successfully classified 98.05% (251/256) of variants assayed, with the remaining being classified as uncertain. We uniformly classified all nonsense variants (14/14) as non-functional and all but one (60/61) synonymous mutation as functional (Figure 3B, C). Missense variants were more often classified as non-functional (121/180) than functional (54/180) (Figure 3B). In total, 44.92% (115/256) of variants were classified as functional and 53.13% (136/256) of variants were classified as non-functional, the remaining 1.95% (5/256) failed to be classified and were designated uncertain.

**Figure 3.**
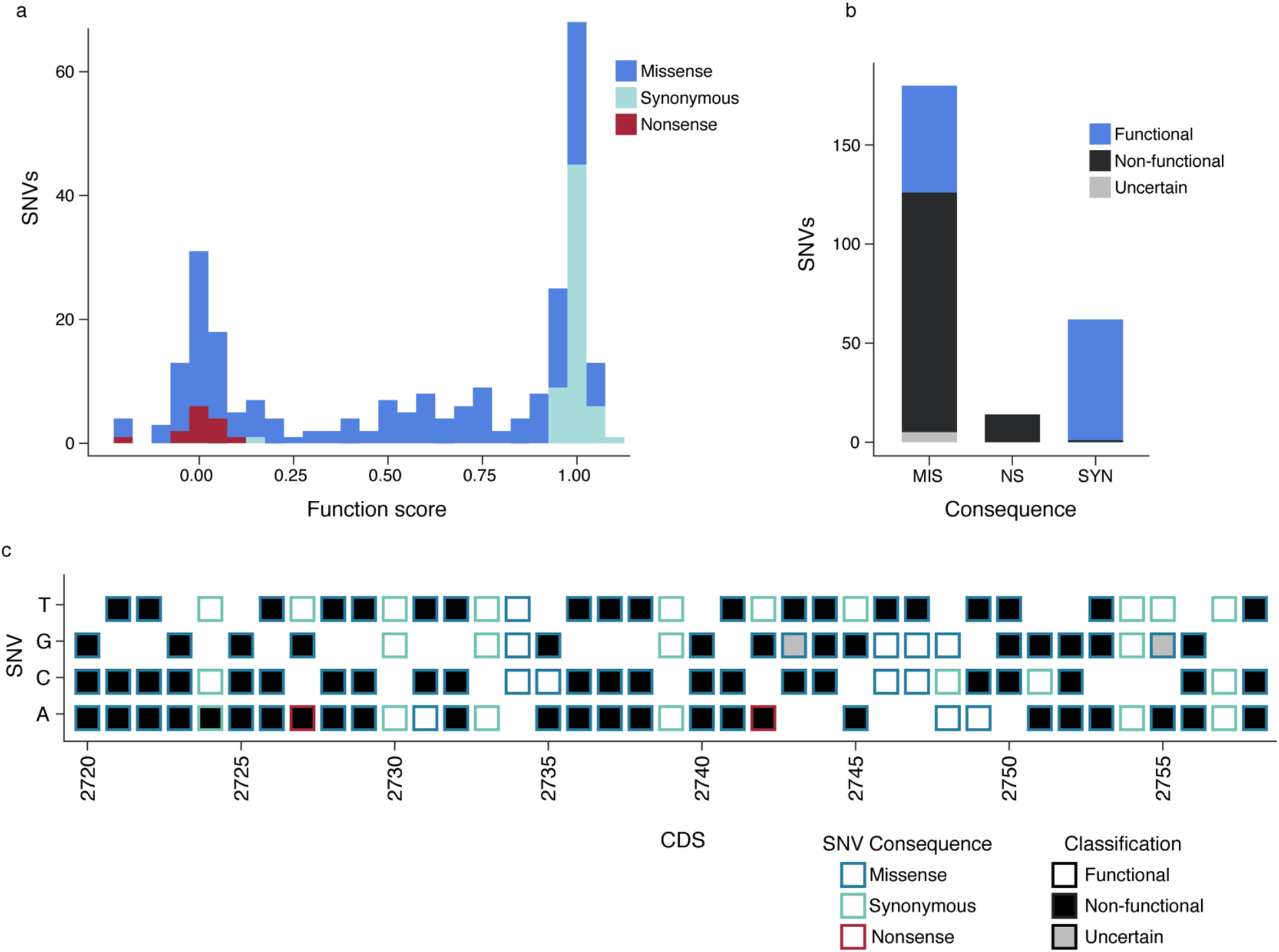
Saturation prime editing enables multiplexed interpretation of *NPC1* variants. (A) Histogram illustrating the distribution of function scores across the 256 variants assayed in *NPC1*, colour indicates mutational consequence. (B) A plot showing the number of SNVs within each mutational category, colour indicates functional classification (MIS, missense; NS, nonsense; SYN, synonymous). (C) A sequence-function map from one of the four saturation prime editing experiments. Boxes are filled according to functional classification and outlined according to mutational consequence. (CDS, coding DNA sequence)

We noted high total editing efficiencies in each library, ranging from 45.56-73.62% (Supplementary Figure S3). In each library, the programmed edits accounted for between 36.53-63.58% of the total sequencing reads, while 5.66-17.19% of the total sequencing reads contained an insertion or deletion or a substitution outside of the targeted area. While each library had a different pegRNA architecture and targeted a different area of *NPC1*, we noted several patterns in the editing outcomes with SPE across the four regions. Similar to previous SGE studies using homology-directed repair^8^, the universal silent PAM-destroying mutation with no secondary mutation was disproportionately the most common editing outcome (Supplementary Figure S4). It has been documented by others that PAM-modifying mutations tend to occur at a greater frequency than other edits made with the same pegRNA in prime editing^11,32^. In our libraries, however, each editing event was programmed to disrupt the PAM sequence directly or indirectly. We speculate that one possible explanation for the preponderance of the PAM-destroying mutation alone could be incomplete reverse-transcription of the pegRNA, leading to partial incorporation of the reverse-transcription template encoding the PAM modifying edit but not the downstream edit-of-interest. In support of this, we found that the three scaffold-proximal nucleotides of the pegRNA tended to be edited with relatively poor efficiency in these multiplexed experiments (Supplementary Figure S4). While detectable in the amplicon sequencing, these edits tended to be represented at such low frequencies that their low read counts made for spurious log-fold changes between replicates. Accordingly, when compiling the scores for classification, we removed edits made by the scaffold-proximal bases, ultimately discarding between 1 to 4 nucleotides or 3 to 12 variants from the downstream analysis.

### Function scores accurately predict pathogenicity and reflect NPC1 protein expression and trafficking

To support the validity of our functional classifications, we first assessed their concordance with definitive clinical interpretations where available. Existing clinical interpretations, however, are relatively sparse for variants in *NPC1*, with only four classifications on ClinVar overlapping with variants assayed in our screens. Encouragingly, the two variants classified as either ‘pathogenic’ or ‘likely pathogenic’ on ClinVar – NPC1 p.G910S and NPC1 p.A1035V – were accurately classified as nonfunctional in our screen while the two variants classified as either ‘benign’ or ‘likely benign’ – NPC1 p.I658I and NPC1 p.V1033V – were both classified as functional. While existing interpretations available for comparison are limited, they do support the accuracy of our functional classifications. Next, we compared our function scores to those provided by the Combined Annotation Dependent Depletion (CADD) computational variant effect prediction software^33^. We found a strong inverse correlation (Spearman’s ρ = −0.74, N = 212) between our function scores and CADD scores (Figure 4A). Notably, the distribution of CADD scores of variants which were classified as functional displayed significant overlap with that of non-functional variants, which tended to have relatively higher CADD scores (Figure 4B). This aligns with the recent findings from SGE-based interpretation of variants in CARD11, where CADD scores appear sensitive, though not highly specific, to non-functional variants^8^.

**Figure 4.**
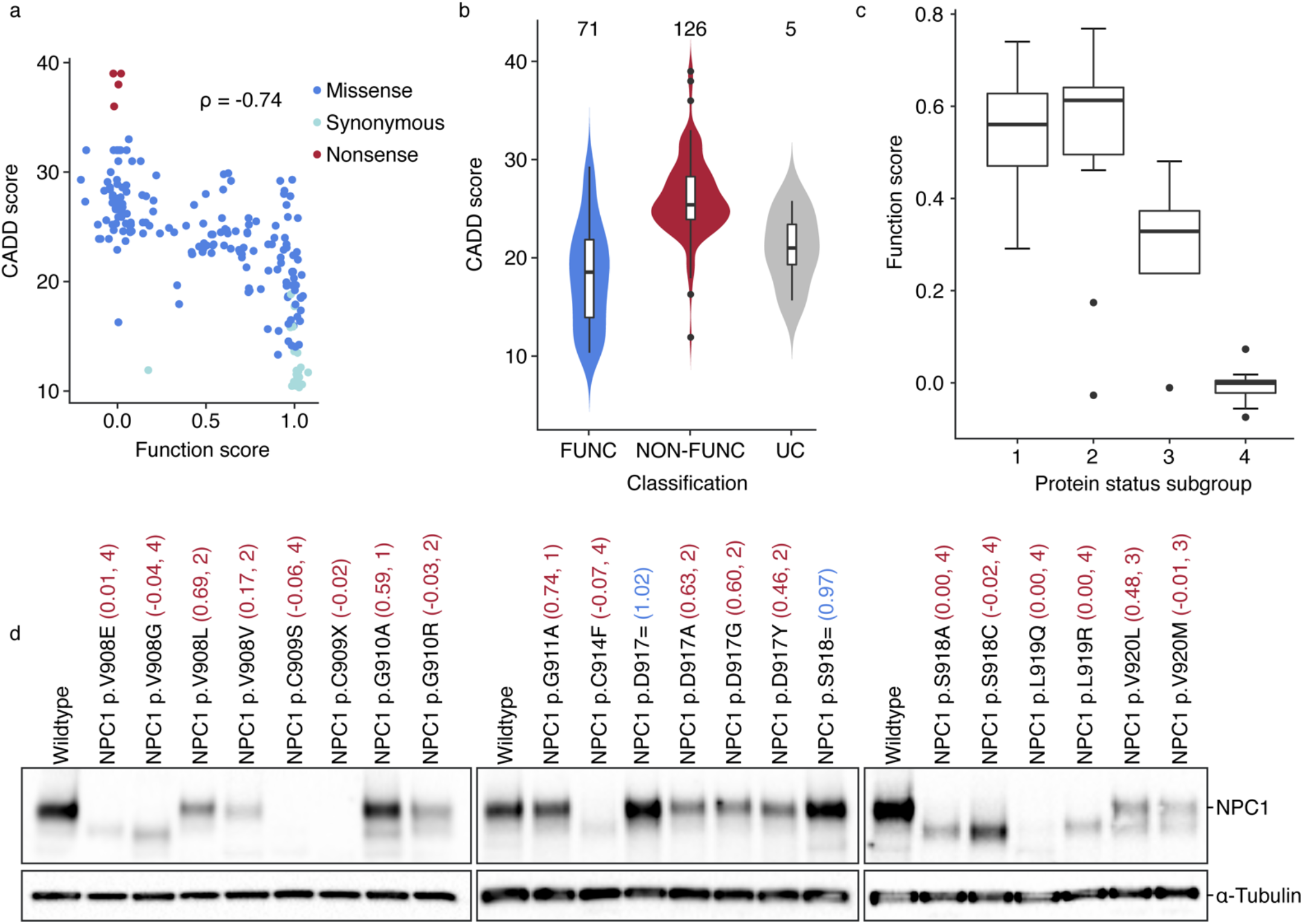
SPE function scores accurately reflect *NPC1* variant pathogenicity. (A) *NPC1* function scores show a strong inverse correlation with CADD scores (Spearman’s ρ = −0.74, N = 212). (B) Violin plots illustrating the distribution of CADD scores within each functional classification. Numbers above indicate number of observations in each group. (FUNC, functional; NON-FUNC, non-functional; UC, uncertain) (C) Distributions of function scores between different protein subgroups. Boxes show the quartiles, whiskers show 1.5 times the interquartile range. (D) Immunoblot for NPC1 from a subset of clones isolated. The function score, followed protein subgroup, is listed in brackets. Colour of function score and protein subgroup indicates functional classification with red being non-functional and blue being functional. Protein subgroups: (1) mutant NPC1 proteins exhibiting similar expression levels and apparent molecular weight compared to wildtype, (2) mutant NPC1 proteins exhibiting decreased protein expression levels but comparable apparent molecular weight with respect to wildtype, (3) mutant NPC1 proteins exhibiting decreased protein expression levels compared to wildtype and two distinct apparent molecular weight products, one matching wildtype and one at a lower molecular weight, and (4) mutant NPC1 proteins exhibiting decreased protein expression levels compared to wildtype and a single lower apparent molecular weight protein product compared to wildtype.

Interestingly, variants which were classified as non-functional in our screens had a wide distribution in function scores. Of note, protein misfolding and incomplete trafficking represent two major molecular disease mechanisms underlying variants in *NPC1*^27,34^. Furthermore, it is well-established that pathogenic missense variants in *NPC1* frequently retain partial functionality^34,35^. Indeed, cholesterol trafficking can be rescued by overexpression of certain *NPC1* mutants, demonstrating that missense mutations do not necessarily ablate protein function^34,35^. It is not well understood, however, what proportion of missense variants are hypomorphic rather than functionally null or if there are varying degrees of partial functionality found between different mutated proteins. Considering this together, we reasoned that the variations found in function scores might reflect differences in protein homeostasis. To address this, we performed preliminary characterization of protein expression and trafficking in 47 different clonal cells lines from our initial SPE screens. We isolated these lines at random by single cell sorting the population of cells after SPE and screening between 12 to 24 clones and expanding all uniquely edited clones. Altogether, this yielded between 4 and 21 unique edited cell clones per targeted region. The NPC1 protein undergoes extensive post-translational modification as it is trafficked from the endoplasmic reticulum to the lysosome and the degree of successful trafficking is reflected in its apparent molecular weight, with mature and immature protein products being distinguishable by western blotting^36^. Using western blotting, we measured NPC1 protein expression and trafficking status in each of the 47 isolated mutants, finding a wide spectrum of defects in both maturation and expression (Supplementary Figure S5). The non-functional mutants with detectable NPC1 protein could be qualitatively classified into four possible subgroups depending on the properties of the detected protein; (1) mutant NPC1 proteins exhibiting similar expression levels and apparent molecular weight compared to wildtype, (2) mutant NPC1 proteins exhibiting decreased protein expression levels but comparable apparent molecular weight with respect to wildtype, (3) mutant NPC1 proteins exhibiting decreased protein expression levels compared to wildtype and two distinct apparent molecular weight products, one matching wildtype and one at a lower molecular weight, and (4) mutant NPC1 proteins exhibiting decreased protein expression levels compared to wildtype and a single lower apparent molecular weight protein product compared to wildtype (Figure 4C, D). We noted that these subgroups were generally well-reflected by the assigned function scores. Specifically, we found that mutants from subgroup 4, which show decreased expression levels and a lower apparent molecular weight compared to wildtype all tend to have function scores similar to the average nonsense function score (Figure 4C). This could indicate that these mutant variants are functionally equivalent to nonsense mutations – functionally null – which aligns with the presumption that the lower molecular weight product is a result of an absence of proper protein maturation and trafficking. Coincident with this, we found that most of the variants with scores that stray from the average nonsense score but are classified as non-functional are represented by mutants which we grouped into categories 1, 2 or 3 (Figure 4C). The exceptions to this are NPC1 p.G910R and NPC1 p.V920M, which have scores similar to the average nonsense score despite having detectable mature protein expression. These results suggest that the differences in function scores assigned in our screens reflect genuine relative functional differences between non-functional variants, despite our binary classification system.

Throughout our screens, we classified one synonymous mutation – NPC1 p.V908V – as non-functional (Figure 3C). Of note, this variant had a function score of 0.17, which is close to the average function score of the nonsense variants assayed. To confirm this classification was accurate, we isolated a clonal NPC1 p.V908V haploidized HEK293T cell clone to characterize the molecular defect. We found that the NPC1 protein in this variant ran at the same molecular weight as a wildtype control indicating proper trafficking was occurring, but there was a marked reduction in total protein expression (Figure 4D). We reasoned that this variant might introduce an errant splicing motif that partially disrupts normal RNA processing. To confirm this, we extracted RNA, reverse transcribed it to cDNA and amplified the edited site from 2 exons up- and downstream of the 18^th^ exon containing this variant. This PCR produced two bands, one corresponding to full-length properly spliced *NPC1* and one shorter product (Supplementary Figure S6). We sequenced both products and while the larger product mapped to normally spliced NPC1 cDNA, the shorter product harboured a 74-base pair deletion. This deletion spanned from two base pairs upstream of the causative G>A mutation to the beginning of the following exon. This suggests the G>A transition underlying the NPC1 p.V908V variant, together with the preceding ‘GT’ dinucleotide, creates a cryptic splice donor which results in the exclusion of the last 74 nucleotides of exon 18. This frameshifting deletion likely results in nonsense mediated decay of the misspliced transcripts, which together with the incomplete penetrance of this variant explain the present, but reduced, NPC1 expression in this cell line. Collectively, these data support our function scores and derived classifications as accurate and clinically relevant. Furthermore, these results indicate that the function scores calculated are useful not only for binary functional classification but could also be used to infer properties about mutant protein expression and trafficking status and ultimately relative functional defects between mutants.

### Extension of saturation prime editing to BRCA2

To demonstrate the versatility of haploidization and saturation prime editing, we applied this variant interpretation approach to the *BRCA2* gene. Deleterious mutations in *BRCA2* significantly increase risk for the development of hereditary breast and ovarian cancers^37^. Fortunately, there are a variety of effective preventative measures available for carriers of these mutations^38,39^. The implementation of these methods, however, is impeded by a poor understanding of the clinical significance of most variants found in the *BRCA2* gene, prioritizing it for SGE.

We modified the haploidization strategy to generate the *BRCA2* haploidized HEK293T cell line by using only one allele-specific guide RNA. Again, using existing wholegenome sequencing data^15^ for the HEK293T cell model we identified a heterozygous polymorphism found just upstream of exon 22 – BRCA2 c.8755-66T>C. The HEK293T model also appeared to be triploid at this locus, with an apparent 2:1 ratio of C:T in both the whole-genome sequencing reads and Sanger sequencing of the locus (Figure 5A). The ‘C’ nucleotide present on two of the alleles completes a potential ‘NGG’ SpCas9 PAM sequence which is absent on the third allele. While the non-targeted allele contained a non-canonical ‘NAG’ SpCas9 PAM^16^, we hypothesized that this would lead to diminished targeting of this allele. Instead of pairing this guide with another heterozygous polymorphism targeting guide RNA, we chose to pair it with a guide RNA that would target all three alleles indiscriminately. We anticipated this would result in a deletion on the two ‘C’-containing alleles, and indel formation at the non-specific guide RNA target-site on the ‘T’-containing allele (Figure 5A). To minimize the influence of a potential indel on *BRCA2* expression or function, we designed the guide RNA to target an intergenic region 4288 base pairs upstream of the start codon of *BRCA2*. We codelivered two plasmid expression vectors each containing one of these two guide RNAs alongside an SpCas9 expression cassette and isolated single-cell clones. We identified a clonal population with the desired deletion product on the two targeted alleles and a 4 base pair deletion at the non-specific target site but no modification at the polymorphic site on the non-targeted allele (Figure 5A). To functionally classify variants in *BRCA2*, we hypothesized that deleterious or loss-of-function variants would elicit a fitness defect discernible over time. To test this, we designed two guide RNAs targeting coding regions within exon 7 and exon 10 and delivered them separately in plasmid expression vectors co-expressing SpCas9 to our *BRCA2* haploidized HEK293T cell. We posited that if loss-of-function variants in *BRCA2* resulted in decreased fitness, initial frameshifting indels formed from CRISPR-Cas9 targeting would deplete over time. Accordingly, we extracted DNA at two timepoints – 4 days post-transfection and 14 days post-transfection – and performed next-generation sequencing of each target site. We found a marked reduction in sequencing reads mapping to frameshifting indels over time, and a corresponding increase in reads mapping to in-frame indels, indicating a fitness defect in cells harbouring frameshifting alleles (Supplementary Figure S7A,B). The frameshifting alleles resulted in premature stop codons that likely ablated *BRCA2* function entirely. To see how deleterious missense variants impact cell fitness, we performed a similar time-course experiment using base editors targeting codons within the DNA binding domain previously determined to be deleterious^40^. We found that sequencing reads which mapped to these missense variants depleted over time in the same manner as a nonsense variant (Supplementary Figure S7C-E). Taken together, these results indicate that deleterious *BRCA2* variants negatively affect cell fitness, and a pooled cell-growth assay could be used to score and classify these variants (Figure 5B).

**Figure 5.**
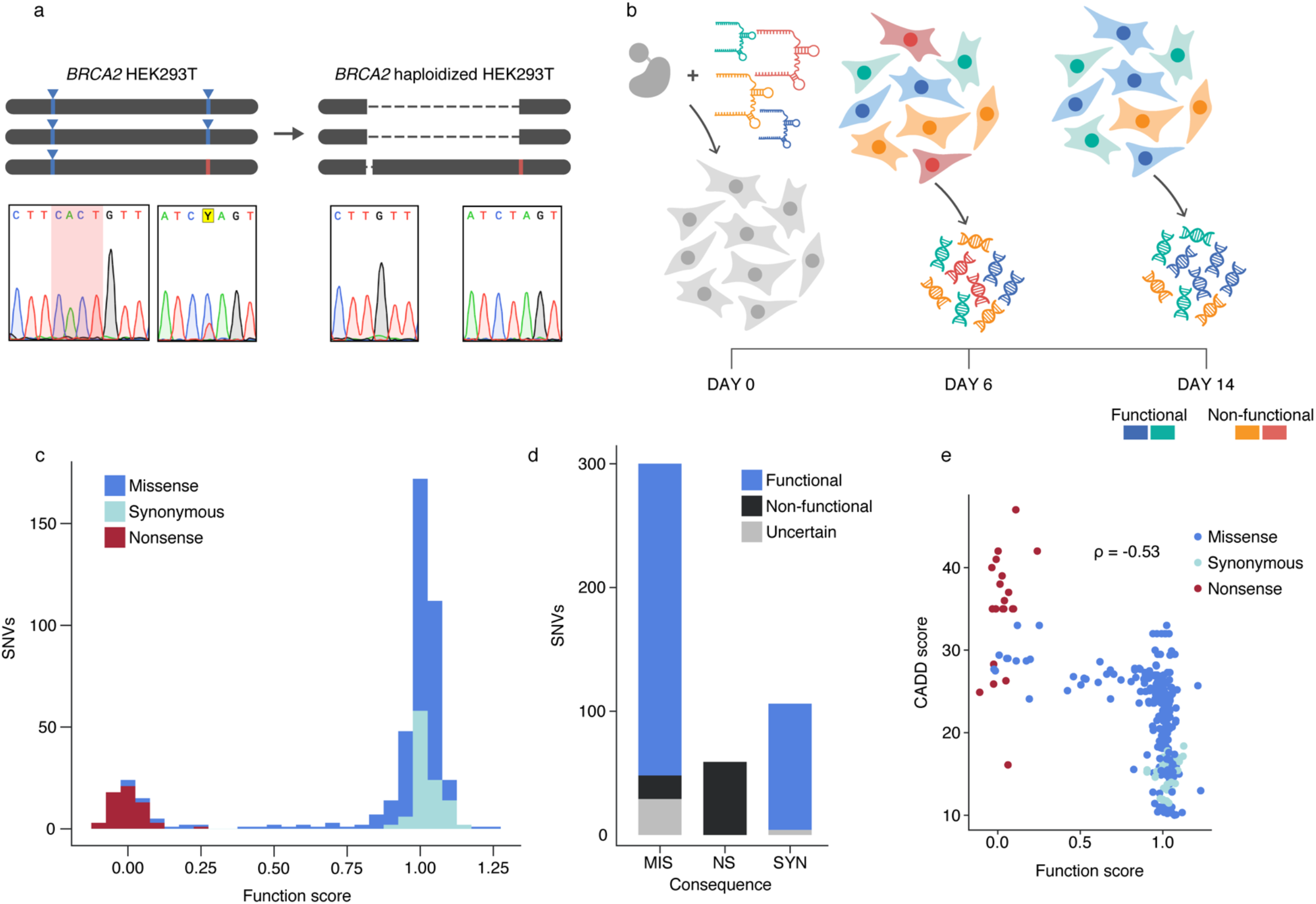
The functional classification of 465 variants in *BRCA2* using SPE. (A) Schematic overview of the haploidization strategy used to generate the *BRCA2* haploidized HEK293T cell line. The nucleotides deleted on the haploidized allele are highlighted in red in the Sanger sequencing read. (B) Schematic illustrating cell growth assay used to classify variants. (C) Histogram illustrating the distribution of function scores across the 465 variants assayed, colour indicates mutational consequence. (D) A plot showing the number of SNVs within each mutational category, colour indicates functional classification (MIS, missense; NS, nonsense; SYN, synonymous). (E) Function scores showed a modest inverse correlation with CADD scores (Spearman’s ρ = −0.53, N = 305).

To perform targeted SPE of the *BRCA2* gene, we selected ten pegRNAs with high computationally predicted editing efficiencies^32^ that spanned the *BRCA2* gene, with a particular focus on the DNA-binding domain. Where possible, we used maximum RT template lengths of 20 nucleotides. Based on our results with targeting *NPC1*, we did not encode mutations in the three pegRNA scaffold-proximal nucleotides, since the editing efficiency at these positions was typically too low to accurately interpret. As with our *NPC1* targeting, we introduced a silent PAM-destroying mutation if the mutation-of-interest did not naturally disrupt the PAM and included pegRNAs encoding additional nonsense mutations by multinucleotide substitutions or by the introduction of a 1-base pair deletion. Altogether, the ten pegRNAs encompassed the following regions: *BRCA2* c.1163-1180, *BRCA2* c.3460-3466, *BRCA2* c.4505-4519, *BRCA2* c.5687-5703, *BRCA2* c.5947-5963, *BRCA2* c.7545-7552, *BRCA2* c.7826-7842, *BRCA2* c.7916-7924 and *BRCA2* c.7930-7960 (Supplementary Table 2). We sampled the population 6 days posttransfection and again at 14 days post-transfection. Functional scoring was accomplished by calculating the log-fold change in read counts for each variant between the two timepoints. We then pooled the scores from each of the ten libraries and scaled their values so that the final average nonsense score was approximately equal to 0 and the silent PAM-destroying allele was approximately equal to 1 (Figure 5C). We performed each editing experiment in 8 replicates and found extremely high reproducibility in the normalized function scores between each replicate (Pearson’s *r* between 0.80 and 0.96, Supplementary Figure S2B). These scores were subsequently used to classify variants as either functional or non-functional using unsupervised clustering (Methods).

In total, we assayed 465 different mutations across the *BRCA2* gene (Figure 5C). We successfully classified 92.90% (432/465) of these variants with 76.12% (354/465) being designated as functional, 16.77% (78/465) being designated as non-functional and the remainder (33/465) inconclusively classified as uncertain. The 59 nonsense mutations assayed were uniformly classified as non-functional and 102 out of the 106 synonymous mutations introduced were classified as functional with the remainder being uncertain. Missense mutations were predominantly functional (252/300) with those classified as non-functional (19/300) all being located within the DNA-binding domain (Figure 5D). Where classifications of ours overlapped with definitive clinical interpretations on ClinVar, we note a near-perfect concordance. The 8 overlapping variants interpreted as ‘likely pathogenic’ or ‘pathogenic’ on ClinVar were classified as non-functional in our assay. Of the 38 overlapping variants interpreted as ‘likely benign’ or ‘benign’ on ClinVar, we classified 36 as functional and 2 as uncertain. The majority (39/47) of variants of uncertain significance on ClinVar that overlapped with our results were classified as functional in our assay, likely due to the proportion of variants queried outside of the DNA binding domain. The remaining 8 variants of uncertain significance were classified as non-functional (3/47) or uncertain (5/47). Of the 7 variants with conflicting interpretations on ClinVar that overlapped with our dataset, 4 were classified as functional, 2 were classified as non-functional, and the remaining variant was classified as uncertain. Similar to *NPC1*, we found a modest inverse correlation (Spearman’s ρ = −0.53, N = 305) between our function scores and CADD scores (Figure 5E). Taken together, these results indicate the broad applicability of cell model haploidization combined with saturation prime editing. The data shown here which indicates the successful translation of this strategy to a second gene with a different functional readout further demonstrates the general utility of this approach.

## Discussion

The present study represents the first demonstration of how CRISPR prime editing can be applied to SGE. We have developed an allele-specific genome editing strategy that produces ‘haploidized’ gene regions, which permits isolated variant analysis. We demonstrated the effectiveness of SPE in these cell lines, providing functional interpretations for 721 SNVs across two different disease-genes using two distinct functional assays.

The allele-specific genome editing strategy utilized is broadly transferable to non-HEK293T cell lines and expands SGE beyond the typical model of choice – HAP1 cells. However, there are caveats worth noting, most notably that this approach is inapplicable to modelling possible dominant-negative effects. Furthermore, if paired heterozygous polymorphisms are used for haploidization, as illustrated in our *NPC1* model, it could be time-consuming to ascertain the phasing of the SNPs-of-interest. The establishment of common haplotypes shared between cell lines, however, could simplify model design. Finally, the establishment of these cell models may be more readily facilitated through the use of ribonucleoprotein (RNP)-based delivery of editing components – which might limit indel formation on the non-targeted allele due to more transient Cas9 exposure.

We note several advantages to SPE. First, we found consistently high editing efficiencies with relatively low indel formation. The modest indel formation we documented could likely be minimized by excluding the nicking guide RNA, though this might diminish total editing efficiency^11^. Notably, we found that editing efficiency was relatively consistent across the targeted area except for the first few pegRNA scaffold-proximal nucleotides. This contrasts with homology-directed repair approaches, where editing efficiency precipitously declines with distance from the double-stranded break^8,41^. These positional biases in editing can require adjustments in the downstream analysis and complicate variant interpretation^7^. Furthermore, all SGE studies to date necessitate the pooling and normalization of scores from different targeted areas across different experiments. In contrast to multiplexed homology-directed repair, it is foreseeable that the prime editing components could be delivered as a single lentiviral library encompassing all SNVs-of-interest, particularly given the highly predictable and pure editing outcomes of prime editing. This would permit the functional interrogation of all SNVs-of-interest in a single experiment, avoiding the need for downstream normalization steps while greatly reducing costs and time-commitment. Lastly, while we focused on SNVs, we did encode multinucleotide edits and single-base pair deletions as part of each experiment. These edits were encoded with similar frequency as the SNVs, indicating that SPE could be expanded to include non-SNV disease-relevant mutations such as small insertions, deletions or duplications.

Several notable limitations of using prime editing over homology-directed repair for SGE include the size of the targetable window and the complexity of target selection. Previously, HDR has been used for SGE of ~100 nucleotide windows^6–8^, more than double our maximum targeted window-size of 43 nucleotides. Our experiments have not, however, defined an absolute upper threshold for targetable window sizes using prime editing and given the high editing efficiencies achieved for this larger window size, we speculate that SPE could support the interrogation of much larger windows sizes dependent on the pegRNA. Furthermore, while the modularity of prime editing – specifically that varying the composition of the 3’-extension can yield different editing outcomes – allows for tunable editing, defining a suitable pegRNA design necessitates thorough optimization. In addition, while mutating the PAM sequence is necessary in all current SGE approaches, this requirement is particularly limiting for SPE, where many highly active pegRNA sequences might be precluded if their PAM is not silently mutable. This could be circumvented by encoding silent mutations in one or in a combination of the three accessible seed-region nucleotides alongside each mutation-of-interest, which might be sufficient to prevent re-binding of the prime editor after an edit has been made. This could also be addressed by using orthogonal Cas9 enzymes with varied PAM specificities, particularly those with longer and thus more flexibly mutable PAM sequences. Encouragingly, the feasibility of prime editing using engineered or orthogonal Cas9 enzymes has recently been demonstrated by multiple groups^42–44^.

In summary, we have developed a generalizable haploidization strategy that simplifies variant interpretation in SGE and showed the applicability of prime editing for saturation variant interpretation. We primarily focused on the *NPC1* gene underlying the rare lysosomal storage disorder Niemann-Pick disease type C1. We demonstrate that the derived function scores are highly reflective of molecular disease mechanisms and could help further our understanding of NPC1 function. For example, we noted several non-functional variants that displayed stable protein expression. While most nonfunctional variants are subject to protein degradation and display varying degrees of trafficking defects, these stably expressed but non-functional variants may highlight residues which are critical to the function of NPC1 as a cholesterol transporter. Furthermore, our function scores show a strong inverse correlation to CADD computational predictions and our functional interpretations are perfectly concordant with the limited existing clinical classifications. To demonstrate the versatility of our experimental design, we extended its demonstration to the highly clinically actionable *BRCA2* gene, functionally classifying 433 different SNVs. We anticipate the described approach will be of immediate utility to others pursuing saturation variant interpretation. Furthermore, the specific function scores and interpretations presented here could help shift currently ambiguous clinical sequencing data into actionable diagnostic information for NPC and *BRCA2*-related hereditary breast and ovarian cancers.

## Methods

### General methods

All genomic DNA isolation was completed using the DNAeasy blood and tissue kit (Qiagen) according to the manufacturers protocol. For genotyping cells and initial estimation of genome editing events, DNA amplification was carried out using DreamTaq polymerase (Thermo Fisher Scientific Scientific). Amplification of products for next-generation sequencing was conducted with Q5 Hot Start High-Fidelity 2X Master Mix (New England BioLabs). All DNA oligonucleotides used in this study were obtained from Integrated DNA Technologies. All plasmids used to express guide RNAs were generated by ligating annealed oligonucleotides into the linearized BPK1520_puroR vector, the assembly of which was described previously^45^. All plasmids used to express pegRNAs, excluding those delivered in libraries, were generated by Golden Gate assembly of annealed oligonucleotides into the pU6-pegRNA-GG-acceptor plasmid, which was a gift from David Liu (Addgene plasmid #132777; http://n2t.net/addgene:132777; RRID: Addgene_132777)^11^. All prime editing experiments were completed using transient expression of the pCMV-PE2 plasmid, which was a gift from David Liu (Addgene plasmid #132775; http://n2t.net/addgene:132775; RRID:Addgene_132775)^11^. Base editing experiments were completed using transient expression of either the pCMV_ABEmax_P2A_GFP vector, which was a gift from David Liu (Addgene plasmid #112101; http://n2t.net/addgene:112101; RRID:Addgene_112101)^46^, the pCAG-CBE4max-SpRY-P2A-EGFP (RTW5133) vector which was a gift from Benjamin Kleinstiver (Addgene plasmid #139999; http://n2t.net/addgene:139999; RRID:Addgene_139999)^47^, or the pCMV-T7-ABEmax(7.10)-SpG-P2A-EGFP (RTW4562) vector, which was a gift from Benjamin Kleinstiver (Addgene plasmid # 140002; http://n2t.net/addgene:140002; RRID:Addgene_140002)^47^. For experiments using an unmodified SpCas9 nuclease, enzyme expression was achieved using transient expression of pSpCas9(BB)-2A-Puro (PX459) V2.0, which was a gift from Feng Zhang (Addgene plasmid #62988; http://n2t.net/addgene:62988; RRID:Addgene_62988)^48^.

### pegRNA design and evaluation

Initial pegRNAs were designed manually. We took the spacer sequences with the highest editing efficiencies predicted by the online software CHOPCHOP (v3)^49^. We manually designed 3’ extension sequences first with 13 nucleotide primer binding sites and reverse transcription templates. We then arbitrarily extended the reverse transcription templates and re-evaluated editing of pegRNAs that produced detectable editing via Sanger sequencing when using the initial 13:13 3’-extension design. Once available, we began using the DeepPE software^32^ for pegRNA design. We found the predictions of DeepPE to be highly accurate, though the longest reverse transcription template length output is 20 nucleotides, limiting its utility for saturation prime editing design. We evaluated each pegRNA by attempting to encode a G>C mutation 6 nucleotides downstream of the pegRNA nick site, thereby mutating the PAM sequence. Outside of saturation prime editing experiments – which were evaluated using next generation amplicon sequencing – we evaluated and quantified all prime editing experiments using Sanger sequencing and the EditR software^50^.

### Polymorphism identification and cell line haploidization

Whole-genome sequencing data from HEK293T cells was accessed from http://hek293genome.org/v2/index.php^15^. Candidate heterozygous polymorphisms were manually identified using the Integrative Genomics Viewer^51^ and their presence was confirmed via Sanger sequencing of genomic DNA. To allow for allele-specific editing, only polymorphisms that generated or destroyed an SpCas9 PAM sequence or occurred within the seed sequence of a potential SpCas9 guide sequence were carried forward for confirmation. Final target selection was based on minimizing off-target sites – determined by the CHOPCHOP v3 software^49^ – and maximizing the portion of the gene removed. Guide RNA sequences were cloned into the PX459 expression vector and transfected in equal proportion into HEK293T cells which then underwent selection and were clonally isolated as described below. For the *NPC1* haploidized cell line, a 15-nucleotide dead guide RNA was designed that placed the non-targeted SNP in the 5^th^ PAM-proximal position. Positive clones were identified by first amplifying the predicted deletion junction formed by the joining of the two cut-sites, then amplifying the two target-sites and sequencing to confirm the sole presence of the non-targeted SNP. To confirm that the deleted fragment was not retained elsewhere in the genome, copy number of the regions both inside and outside the targeted space was quantified by the Centre for Applied Genomics, the Hospital for Sick Children, Toronto, Ontario using digital droplet PCR. In addition, each cell clone was genotyped at each position where a heterozygous SNP was present in the parental HEK293T cell line to confirm complete haploidization of the region-of-interest.

### General cell culture conditions

HEK293T cells were cultured in Dulbecco’s Modified Eagle’s Medium (Wisent) supplemented with 10% fetal bovine serum (Wisent) and 1% penicillin streptomycin (Wisent) and incubated and maintained in a humidified chamber at 37C with 5% CO2. The HEK293T cell line used in this study was authenticated by short tandem repeat analysis performed by the Centre for Applied Genomics, the Hospital for Sick Children, Toronto, Ontario.

### Protein isolation and western blotting

Cells were trypsinized, pelleted, and washed three times with 1× PBS (Wisent). A protein extraction buffer consisting of a one-to-one solution of RIPA homogenizing buffer (50 mM Tris HCl at pH 7.4, 150 nM NaCl, 1-mM EDTA) and RIPA double-detergent buffer (2% deoxycholate, 2% NP-40, 2% Triton X-100 in RIPA homogenizing buffer) supplemented with a protease-inhibitor cocktail (Roche) was used to resuspend the cells, which were then incubated on ice for 30 minutes. After centrifuging the cells at 12,000g for 15 minutes at 4°C, the supernatant was collected and stored at −80°C. The protein concentration of the lysate was measured using the Pierce BCA protein assay kit according to the manufacturer’s protocol (Thermo Fisher Scientific Scientific). SDS-Page separation was completed by running between 3 and 10μg of total protein on a NuPAGE 3%–8% Tris-acetate gel (Thermo Fisher Scientific Scientific). Next, protein transfer to a nitrocellulose membrane was completed using the iBlot 2 Dry Blotting System (Thermo Fisher Scientific Scientific). The membrane was then blocked for one hour at room temperature using a 5% milk solution in 1× TBST. Next, the membrane was then incubated in NPC1 (Abcam, ab106534) and α-Tubulin (Sigma Aldrich, T6199) primary antibodies overnight at 4°C. The next day, the primary antibody solution was removed, and the membrane was washed three times with 1× TBST. This was followed by incubation horseradish peroxidase conjugated goat anti-rabbit IgG (Abcam, ab6721) for NPC1 and anti-mouse IgG (Abcam, ab205719) for α-Tubulin for 1 hour at room temperature. After three washes with 1× TBST, signal was collected using the SuperSignal West Femto Maximum Sensitivity Substrate (Thermo Fisher Scientific Scientific) according to the manufacturer’s protocol.

### Fluorescent staining and imaging

One day before staining, 200,000 cells we seeded on glass coverslips in a 12-well plate. The next day, cells were stained for 1 hour with 2μm of LysoTracker™ Red DND-99 (Thermo Fisher Scientific), then fixed with 4% paraformaldehyde in 1× PBS (Wisent). Following fixation, nuclear staining with Hoechst 34580 (Thermo Fisher Scientific) was carried out over a 15-minute incubation period. Finally, cells were washed and mounted onto slides using ProLong Gold Antifade Mountant (Thermo Fisher Scientific). Slides were visualized using a Quorum spinning disk confocal microscope (Quorum Technologies) with images captured using a Hamamatsu C9100-13 EM-CCD camera.

### Cell transfection

All transfections were carried out by first seeding cells at 600,000 cells per well on 6-well plates. For base editing experiments, 1000ng of the base editor expression plasmid was co-transfected with 1000ng the guide RNA expression plasmid. For experiments using an unmodified SpCas9 nuclease, 1000ng of the SpCas9 expression plasmid was co-transfected with 1000ng of the guide RNA expression plasmid. For all experiments involving prime editing, 700ng of the pegRNA expression plasmid, 500ng of the prime editor expression plasmid and 250ng of the nicking guide RNA expression plasmid were co-transfected. Twelve to twenty-four hours post-transfection, puromycin was added to the cells at 2μg/mL in complete medium. This selection medium was replaced every 24 hours for the following 3 days then switched back to puromycin-free complete medium until the relevant experiments were completed.

### Cell line isolation

HEK293T cells were seeded at 250,000 cells per well on 12-well plates (Corning). The following day, cells were transfected as described above. 24 hours post-transfection, puromycin (Gibco) was added at a concentration of 2μg/mL in complete medium and changed daily for the following 3 days. Next, cells were trypsinized and resuspended in FACS buffer composed of PBS without calcium or magnesium, supplemented with 2% FBS and 2.5 mM EDTA, at a concentration of 1,000,000 cells per mL. Cells were then passed through a 40 μm cell strainer and then sorted using a MoFloXDP cell sorter (Beckman Coulter) into a 96-well plate containing complete medium. Clonal lines were expanded for 14 days before passaged for genomic DNA extraction and genotyping.

### Prime editing guide RNA library design and cloning

For each targeted site, an assembly vector was generated from the pU6-pegRNA-GG-acceptor plasmid. First, the full-length pegRNA was cloned into this vector by Golden Gate assembly. Next, InFusion cloning (Takara) was used to replace the reverse transcription template portion of the 3’-extension with the BsaI-flanked red-fluorescent protein cassette from the original pU6-pegRNA-GG-acceptor plasmid. This construct was then digested with BsaI to drop out the red-fluorescent protein cassette and the resulting linearized vector was used for library assembly. To generate the final library of pegRNAs, an oPool Oligo Pool (Integrated DNA Technologies) was synthesized containing each possible reverse-transcription template necessary to effectuate saturational mutagenesis of the targeted region flanked by 30 nucleotides of homology to the destination vector. This single-stranded oligo pool was then combined with the linearized library vector for assembly using the NEBuilder® HiFi DNA Assembly Master Mix (New England BioLabs) according to the manufacturer’s protocol. The assembly reaction was then transformed into Stellar competent cells (Takara), and following an hour-long outgrowth, the transformed cells were directly transferred into 300mL of ampicillin-containing media for 24 hours. The final plasmid library DNA was isolated from the culture using the Nucleobond Xtra Maxi kit (Macherey-Nagel).

### Pooled functional assay for NPC1 variants

For transfection of pegRNA libraries, *NPC1* haploidized HEK293T cells were seeded at 600,000 cells per well on 6-well plates. Each of the 6-wells were transfected with 700 ng of the pegRNA library plasmid pool, 500 ng of pCMV-PE2, and 250 ng of BPK1520_puroR containing the relevant nicking guide RNA. After three days of selection, we pooled the cells into a flask and allowed cells to expand for seven days. Next, cells were passaged and seeded at 2,000,000 cells on 60 mm plates in at least 8 replicates. The following day, cells were stained with 2 μm of LysoTracker™ Green DND-26 (Thermo Fisher Scientific) in DMEM for 1 hour, then subject to FACS using a Sony SH800S Cell Sorter (Sony). We first sorted stained unedited control NPC1 haploidized HEK293T cells to define the sorting gates. We defined two sorting gates each accounting for ~ 50% of the population: one ‘low’ gate encompassing the dimmest 50% of fluorescent cells and one ‘high’ gate encompassing the brightest 50% of fluorescent cells. Each of the experimental replicates were sorted using these gates, collecting a minimum of 300,000 cells per gate. We then extracted genomic DNA from each subpopulation and amplified the targeted region with primers flanked by universal Illumina adaptors for subsequent library preparation and sequencing.

### Pooled functional assay for BRCA2 variants

Initial assay development was completed using unmodified SpCas9 to generate indels within the coding region of *BRCA2*. We extracted genomic DNA 4 days and 14 days post-transfection, then amplified and used high-throughput sequencing to quantify the editing events. We then performed a similar time course experiment using base editors which targeted three distinct areas of the DNA-binding domain. In these experiments, DNA was extracted 6 days and 16 days post-transfection with editing events quantified using high-throughput sequencing.

For transfection of pegRNA libraries in our saturation prime editing screens, haploidized HEK293T cells were seeded at 1,00,000 cells per well on 60mm plates in 8 replicates. Each of the plates were transfected with 700ng of the pegRNA library plasmid pool, 500ng of pCMV-PE2, and 250ng of BPK1520_puroR containing the relevant nicking guide RNA. Transfected cells were selected with 2*μ*g/mL of puromycin for 3 days then allowed to recover for 3 more days. At this point, cells were washed and passaged, and half the population of each replicate was reseeded in a 60mm plate to expand for an additional 10 days while the other half was used for genomic DNA extraction, representing the first timepoint. The expanding cells were passaged every 2-days, reseeding 1/3^rd^ of the population with each passage. On the 16^th^ day post-transfection, all remaining cells were collected for genomic DNA extraction, representing the final timepoint. Finally, the targeted region was amplified with primers flanked by universal Illumina adaptors for subsequent library preparation and sequencing.

### Nucleic acid sampling and sequencing

In both *NPC1* and *BRCA2* experiments, all genomic DNA extracted was used for subsequent amplification, with 1000ng being used per reaction in multiple simultaneous PCR reactions. This initial amplification appended universal Illumina adaptor sequences that served to prime for indexing library preparation. To avoid spurious perturbation in variant distribution, PCR reactions were carried out with minimal cycling. All reactions were pooled and purified using the QIAquick PCR purification Kit (Qiagen) following the manufacturer’s protocol. These were then submitted for Indexing PCR to the Donnelly Sequencing Centre at the University of Toronto (http://ccbr.utoronto.ca/donnelly-sequencing-centre). Samples were first quantified with fluorescent chemistry in the Quant-iT 1X dsDNA HS Assay (Thermo Fisher Scientific) then 10 ng per sample was processed using HotStarTaq Plus Master Mix Kit (Qiagen), with 5 cycles of amplification. Illumina’s non-redundant unique dual indexes (UDI) were used in this step (IDT for Illumina Nextera DNA Unique Dual Indexes Set C, Illumina Inc.). Indexed PCR amplicons were purified using HighPrep PCR Clean-up magnetic beads (MagBio Genomics) at ratio of 0.9:1 (beads:sample). Next, indexed purified PCR amplicons were quantified using the Quant-iT 1X dsDNA HS Assay (Thermo Fisher Scientific), and were pooled at equimolar ratios after size-adjustment. The final pool was run on an Agilent Bioanalyzer dsDNA High Sensitivity chip (Agilent Technologies Inc.) and quantified using NEBNext Library Quant Kit for Illumina (New England Biolabs). The quantified pool was hybridized at a final concentration of 2.2 pM and sequenced paired-end on the Illumina NextSeq 500 platform using a High-Output v2.5 flowcell and 2×151c read length.

### Data analysis and variant scoring

Next-generation sequencing reads were processed using the CRISPResso2 software (version 2.0.40) using the default parameters, except for experiments involving base editors where base editor mode was selected. The read counts for each allele found in output file “Alleles_frequency_table_around_sgRNA” from CRISPResso2 were used to calculate changes across timepoints in the case of *BRCA2* experiments, and across gated populations in the case of *NPC1* experiments. For analyzing the distribution of indel sizes over time when developing the BRCA2 functional assay, alleles with fewer than 100 reads were filtered from analysis.

For *NPC1*, function scores were derived by first calculating the log-fold change from the low-to high-fluorescent population then the log-fold change value of the wildtype sequencing read was subtracted from each score. Next, each score was divided by the average score of the nonsense mutations assayed. Finally, scores were multiplied by −1, then increased by 1 such that the average nonsense score was approximately 0, and the average function score was approximately 1. For *BRCA2*, we first calculated the logfold change of each allele from the second to first timepoint. We noted that the wildtype allele depleted from the first to second timepoint, indicating the prime editing had not reached completion at our first timepoint. Accordingly, to account for continued editing over our experiment we centered our scores on the silent PAM-destroying mutation, subtracting its log-fold change value from each score. Like *NPC1*, we then divided each score by the average score of the nonsense mutations assayed. The final function score was arrived at by then multiplying each score by −1 and then increasing it by 1.

Rather than choosing cutoff scores for classification, we elected to take a more data driven approach. We observed that the benign cluster tends to be large and tightly bounded, and while there is a noticeable nonsense cluster, non-functional variants tend to display a wide variance. Our classification is based on defining what scores constitute the functional cluster, then classifying everything with a score below the lower bound of this cluster as functional. To define the functional cluster, we use unsupervised meanshift clustering^52^. Specifically, we seed a single cluster with a score of 1 – the centering adjustments to our scores ensures the cluster center should be approximately equal to 1 – and then implement the mean-shift algorithm using Python’s scikit-learn package^53^. All function scores that fall within this cluster will have their corresponding variant classified as functional. Arriving at the function score and this classification, however, required several processing steps – centering the data such that the wildtype or PAM-destroying mutation was approximately equal to 1 and the average nonsense score was approximately equal to 0, normalizing the data across different experiments and the mean-shift clustering – each of which might introduce some degree of error. To account for this, we adopted a resampling-based approach. Specifically, we resample by taking a permutation of the eight experimental replicates and repeat the entire data analysis pipeline – centering, normalizing, and classifying – and repeat this 1000 times. We only retain classifications if they were consistent across at least 998/1000 resampling with anything not meeting this cutoff being classified as uncertain.

## External data sources

Variant annotations from CADD (version 1.5) and ClinVar (Downloaded 27 January 2020) were provided as part of the variant annotation pipeline from the Centre for Applied Genomics, The Hospital for Sick Children, Toronto, Ontario. Coding nucleotide and amino acid positions reported in this study correspond to the NCBI transcript NM_000271 for *NPC1* and NM_000059 for *BRCA2*.

## Data availability

Next-generation sequencing data have been deposited to the NCBI Sequence Read Archive database under accession PRJNA728726.

## Acknowledgements

This study was supported by Niemann-Pick Canada and the SickKids Foundation. Additional support was provided by a research grant from the University of Pennsylvania Orphan Disease Center in partnership with The Andrew Coppola Foundation (MDBR-21-113-NPC to E.A.I). We also thank the Hospital for Sick Children (Restracomp to S.E). We wish to thank Bhooma Thiruvahindrapuram and Thomas Nalpathamkalam of the Centre for Applied Genomics, The Hospital for Sick Children, Toronto, Canada for assistance with applying their variant annotation pipeline. We are grateful to Tanja Durbic and the rest of the Donnelly Sequencing Centre staff for their assistance with sequencing experiments.

## References

1 Cooper, G. M. Parlez-vous VUS? Genome Res 25, 1423–1426, doi:10.1101/gr.190116.115 (2015).

2 Starita, L. M. et al. Variant Interpretation: Functional Assays to the Rescue. Am J Hum Genet 101, 315–325, doi:10.1016/j.ajhg.2017.07.014 (2017).

3 Fowler, D. M. & Fields, S. Deep mutational scanning: a new style of protein science. Nat Methods 11, 801–807, doi:10.1038/nmeth.3027 (2014).

4 Majithia, A. R. et al. Prospective functional classification of all possible missense variants in PPARG. Nat Genet 48, 1570–1575, doi:10.1038/ng.3700 (2016).

5 Sun, S. et al. A proactive genotype-to-patient-phenotype map for cystathionine beta-synthase. Genome Med 12, 13, doi:10.1186/s13073-020-0711-1 (2020).

6 Findlay, G. M., Boyle, E. A., Hause, R. J., Klein, J. C. & Shendure, J. Saturation editing of genomic regions by multiplex homology-directed repair. Nature 513, 120–123, doi:10.1038/nature13695 (2014).

7 Findlay, G. M. et al. Accurate classification of BRCA1 variants with saturation genome editing. Nature 562, 217–222, doi:10.1038/s41586-018-0461-z (2018).

8 Meitlis, I. et al. Multiplexed Functional Assessment of Genetic Variants in CARD11. Am J Hum Genet 107, 1029–1043, doi:10.1016/j.ajhg.2020.10.015 (2020).

9 Kweon, J. et al. A CRISPR-based base-editing screen for the functional assessment of BRCA1 variants. Oncogene 39, 30–35, doi:10.1038/s41388-019-0968-2 (2020).

10 Hanna, R. E. et al. Massively parallel assessment of human variants with base editor screens. Cell 184, 1064–1080 e1020, doi:10.1016/j.cell.2021.01.012 (2021).

11 Anzalone, A. V. et al. Search-and-replace genome editing without double-strand breaks or donor DNA. Nature 576, 149–157, doi:10.1038/s41586-019-1711-4 (2019).

12 Carette, J. E. et al. Haploid genetic screens in human cells identify host factors used by pathogens. Science 326, 1231–1235, doi:10.1126/science.1178955 (2009).

13 Vanier, M. T. Niemann-Pick disease type C. Orphanet J Rare Dis 5, 16, doi:10.1186/1750-1172-5-16 (2010).

14 Wassif, C. A. et al. High incidence of unrecognized visceral/neurological late-onset Niemann-Pick disease, type C1, predicted by analysis of massively parallel sequencing data sets. Genet Med 18, 41–48, doi:10.1038/gim.2015.25 (2016).

15 Lin, Y. C. et al. Genome dynamics of the human embryonic kidney 293 lineage in response to cell biology manipulations. Nat Commun 5, 4767, doi:10.1038/ncomms5767 (2014).

16 Hsu, P. D. et al. DNA targeting specificity of RNA-guided Cas9 nucleases. Nat Biotechnol 31, 827–832, doi:10.1038/nbt.2647 (2013).

17 Cong, L. et al. Multiplex genome engineering using CRISPR/Cas systems. Science 339, 819–823, doi:10.1126/science.1231143 (2013).

18 Wu, X. et al. Genome-wide binding of the CRISPR endonuclease Cas9 in mammalian cells. Nat Biotechnol 32, 670–676, doi:10.1038/nbt.2889 (2014).

19 Duan, J. et al. Genome-wide identification of CRISPR/Cas9 off-targets in human genome. Cell Res 24, 1009–1012, doi:10.1038/cr.2014.87 (2014).

20 Rose, J. C. et al. Suppression of unwanted CRISPR-Cas9 editing by coadministration of catalytically inactivating truncated guide RNAs. Nat Commun 11, 2697, doi:10.1038/s41467-020-16542-9 (2020).

21 Coelho, M. A. et al. CRISPR GUARD protects off-target sites from Cas9 nuclease activity using short guide RNAs. Nat Commun 11, 4132, doi:10.1038/s41467-020-17952-5 (2020).

22 Kopitz, J., Gerhard, C., Hofler, P. & Cantz, M. [14C]Methylamine accumulation in cultured human skin fibroblasts--a biochemical test for lysosomal storage and lysosomal diseases. Clin Chim Acta 227, 121–133, doi:10.1016/0009-8981(94)90141-4 (1994).

23 Lachmann, R. H. et al. Treatment with miglustat reverses the lipid-trafficking defect in Niemann-Pick disease type C. Neurobiol Dis 16, 654–658, doi:10.1016/j.nbd.2004.05.002 (2004).

24 te Vruchte, D. et al. Relative acidic compartment volume as a lysosomal storage disorder-associated biomarker. J Clin Invest 124, 1320–1328, doi:10.1172/JCI72835 (2014).

25 Xu, M. et al. A phenotypic compound screening assay for lysosomal storage diseases. J Biomol Screen 19, 168–175, doi:10.1177/1087057113501197 (2014).

26 Millat, G. et al. Niemann-Pick C1 disease: the I1061T substitution is a frequent mutant allele in patients of Western European descent and correlates with a classic juvenile phenotype. Am J Hum Genet 65, 1321–1329, doi:10.1086/302626 (1999).

27 Millat, G. et al. Niemann-Pick C1 disease: correlations between NPC1 mutations, levels of NPC1 protein, and phenotypes emphasize the functional significance of the putative sterol-sensing domain and of the cysteine-rich luminal loop. Am J Hum Genet 68, 1373–1385, doi:10.1086/320606 (2001).

28 Dardis, A. et al. Molecular Genetics of Niemann-Pick Type C Disease in Italy: An Update on 105 Patients and Description of 18 NPC1 Novel Variants. J Clin Med 9, doi:10.3390/jcm9030679 (2020).

29 Millat, G. et al. Niemann-Pick C disease: use of denaturing high performance liquid chromatography for the detection of NPC1 and NPC2 genetic variations and impact on management of patients and families. Mol Genet Metab 86, 220–232, doi:10.1016/j.ymgme.2005.07.007 (2005).

30 Greer, W. L. et al. Mutations in NPC1 highlight a conserved NPC1-specific cysteine-rich domain. Am J Hum Genet 65, 1252–1260, doi:10.1086/302620 (1999).

31 Anzalone, A. V., Koblan, L. W. & Liu, D. R. Genome editing with CRISPR-Cas nucleases, base editors, transposases and prime editors. Nat Biotechnol 38, 824–844, doi:10.1038/s41587-020-0561-9 (2020).

32 Kim, H. K. et al. Predicting the efficiency of prime editing guide RNAs in human cells. Nat Biotechnol 39, 198–206, doi:10.1038/s41587-020-0677-y (2021).

33 Rentzsch, P., Witten, D., Cooper, G. M., Shendure, J. & Kircher, M. CADD: predicting the deleteriousness of variants throughout the human genome. Nucleic Acids Res 47, D886–D894, doi:10.1093/nar/gky1016 (2019).

34 Gelsthorpe, M. E. et al. Niemann-Pick type C1 I1061T mutant encodes a functional protein that is selected for endoplasmic reticulum-associated degradation due to protein misfolding. J Biol Chem 283, 8229–8236, doi:10.1074/jbc.M708735200 (2008).

35 Zampieri, S., Bembi, B., Rosso, N., Filocamo, M. & Dardis, A. Treatment of Human Fibroblasts Carrying NPC1 Missense Mutations with MG132 Leads to an Improvement of Intracellular Cholesterol Trafficking. JIMD Rep 2, 59–69, doi:10.1007/8904_2011_49 (2012).

36 Watari, H. et al. Mutations in the leucine zipper motif and sterol-sensing domain inactivate the Niemann-Pick C1 glycoprotein. J Biol Chem 274, 21861–21866, doi:10.1074/jbc.274.31.21861 (1999).

37 Kuchenbaecker, K. B. et al. Risks of Breast, Ovarian, and Contralateral Breast Cancer for BRCA1 and BRCA2 Mutation Carriers. JAMA 317, 2402–2416, doi:10.1001/jama.2017.7112 (2017).

38 Olopade, O. I. & Artioli, G. Efficacy of risk-reducing salpingo-oophorectomy in women with BRCA-1 and BRCA-2 mutations. Breast J 10 Suppl 1, S5–9, doi:10.1111/j.1524-4741.2004.101s3.x (2004).

39 Rebbeck, T. R. et al. Bilateral prophylactic mastectomy reduces breast cancer risk in BRCA1 and BRCA2 mutation carriers: the PROSE Study Group. J Clin Oncol 22, 1055–1062, doi:10.1200/JCO.2004.04.188 (2004).

40 Hart, S. N. et al. Comprehensive annotation of BRCA1 and BRCA2 missense variants by functionally validated sequence-based computational prediction models. Genet Med 21, 71–80, doi:10.1038/s41436-018-0018-4 (2019).

41 Paquet, D. et al. Efficient introduction of specific homozygous and heterozygous mutations using CRISPR/Cas9. Nature 533, 125–129, doi:10.1038/nature17664 (2016).

42 Kweon, J. et al. Engineered prime editors with PAM flexibility. Mol Ther, doi:10.1016/j.ymthe.2021.02.022 (2021).

43 Liu, P. et al. Improved prime editors enable pathogenic allele correction and cancer modelling in adult mice. Nat Commun 12, 2121, doi:10.1038/s41467-021-22295-w (2021).

44 Hua, K., Jiang, Y., Tao, X. & Zhu, J. K. Precision genome engineering in rice using prime editing system. Plant Biotechnol J 18, 2167–2169, doi:10.1111/pbi.13395 (2020).

45 Erwood, S. et al. Modeling Niemann-Pick disease type C in a human haploid cell line allows for patient variant characterization and clinical interpretation. Genome Res 29, 2010–2019, doi:10.1101/gr.250720.119 (2019).

46 Koblan, L. W. et al. Improving cytidine and adenine base editors by expression optimization and ancestral reconstruction. Nat Biotechnol 36, 843–846, doi:10.1038/nbt.4172 (2018).

47 Walton, R. T., Christie, K. A., Whittaker, M. N. & Kleinstiver, B. P. Unconstrained genome targeting with near-PAMless engineered CRISPR-Cas9 variants. Science 368, 290–296, doi:10.1126/science.aba8853 (2020).

48 Ran, F. A. et al. Genome engineering using the CRISPR-Cas9 system. Nat Protoc 8, 2281–2308, doi:10.1038/nprot.2013.143 (2013).

49 Labun, K. et al. CHOPCHOP v3: expanding the CRISPR web toolbox beyond genome editing. Nucleic Acids Res 47, W171–W174, doi:10.1093/nar/gkz365 (2019).

50 Kluesner, M. G. et al. EditR: A Method to Quantify Base Editing from Sanger Sequencing. CRISPR J 1, 239–250, doi:10.1089/crispr.2018.0014 (2018).

51 Robinson, J. T. et al. Integrative genomics viewer. Nat Biotechnol 29, 24–26, doi:10.1038/nbt.1754 (2011).

52 Comaniciu, D. & Meer, P. Mean shift: a robust approach toward feature space analysis. IEEE Transactions on Pattern Analysis and Machine Intelligence 24, 603–619, doi:10.1109/34.1000236 (2002).

53 Pedregosa, F. et al. Scikit-learn: Machine Learning in Python. J. Mach. Learn. Res. 12, 2825–2830 (2011).

