## Supplementary figures and images for "Saturation variant interpretation using CRISPR prime editing"

### Supplementary Figure S1

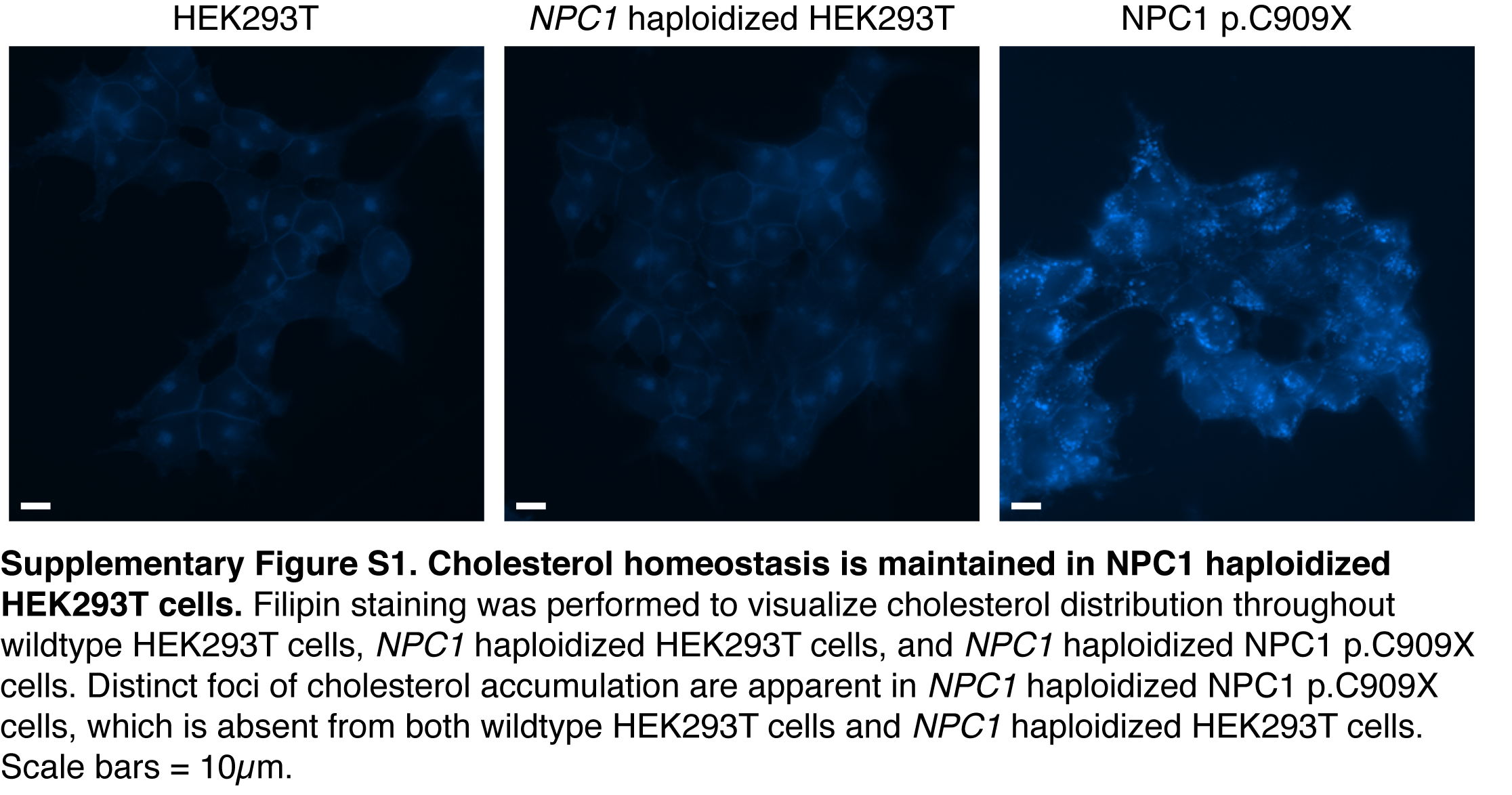

### Supplementary Figure S2

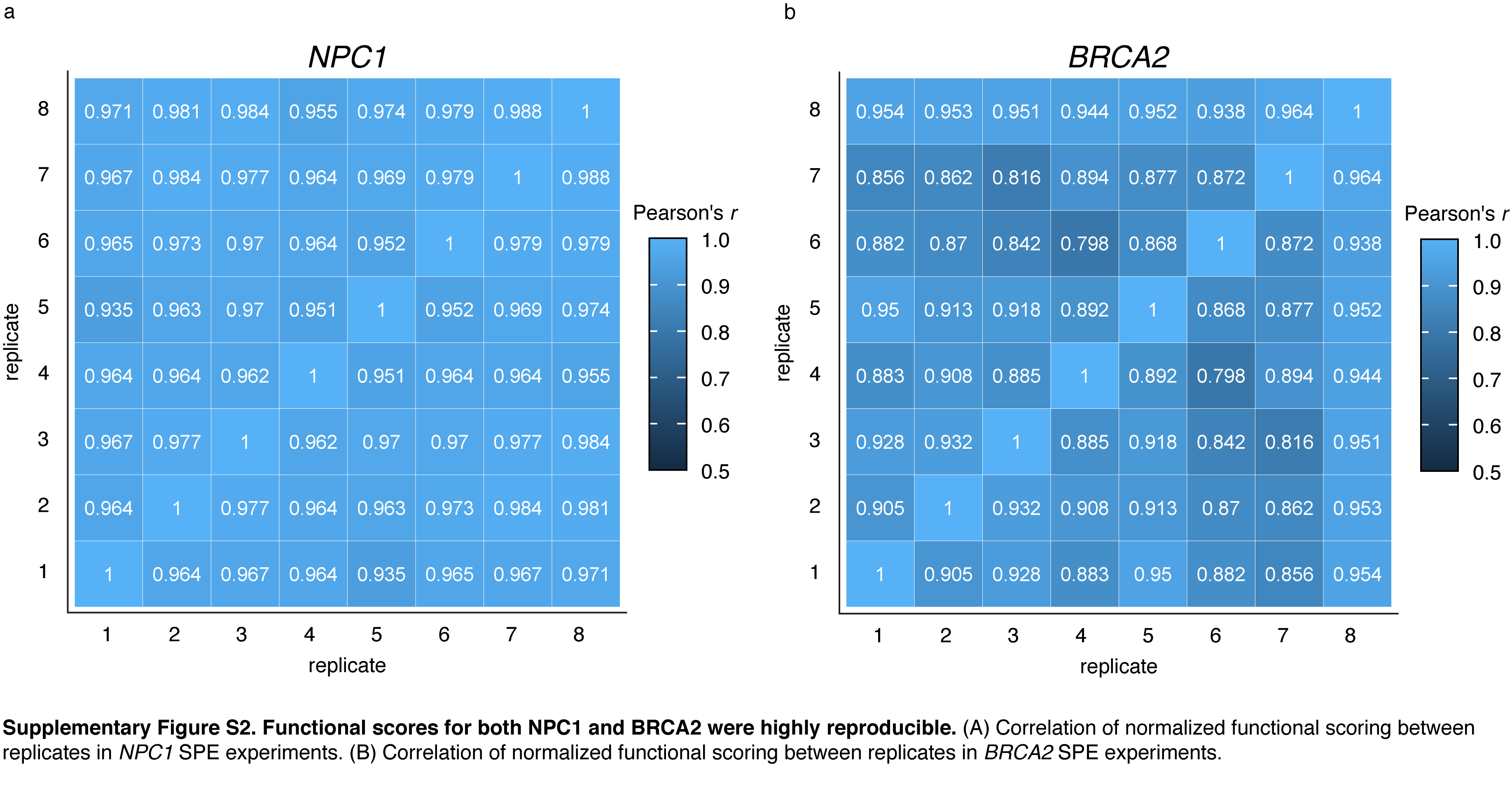

### Supplementary Figure S3

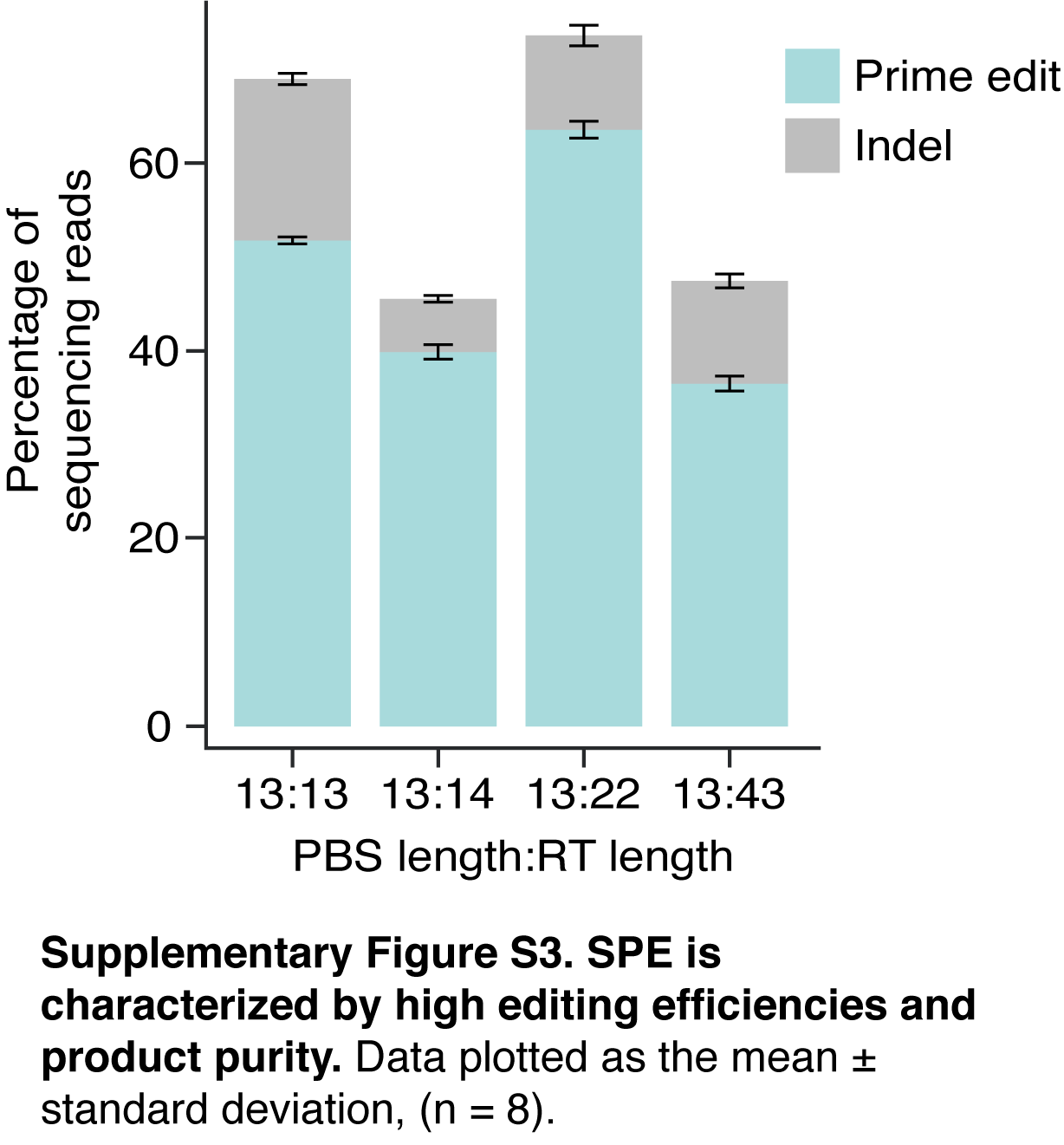

### Supplementary Figure S4

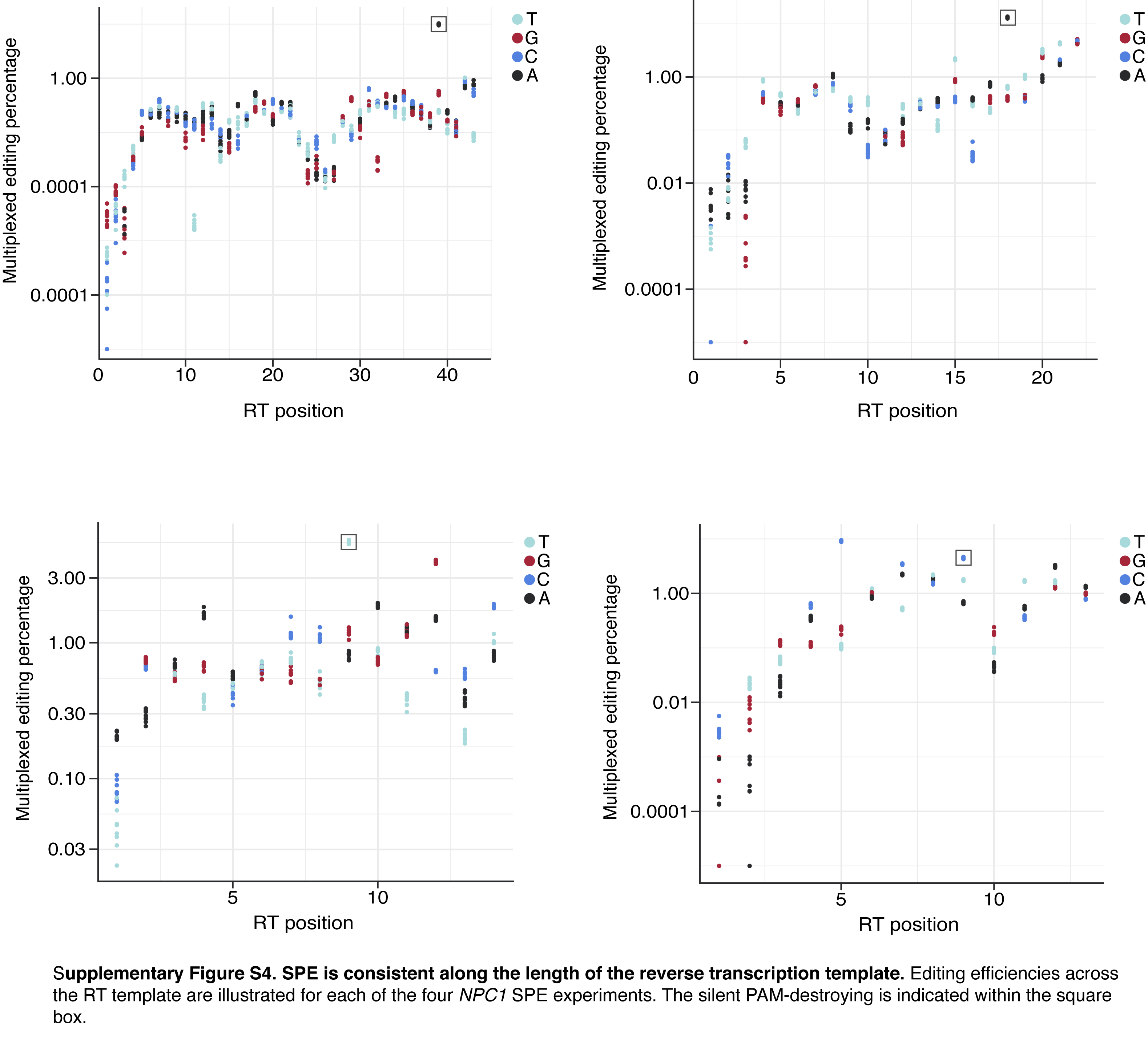

### Supplementary Figure S5

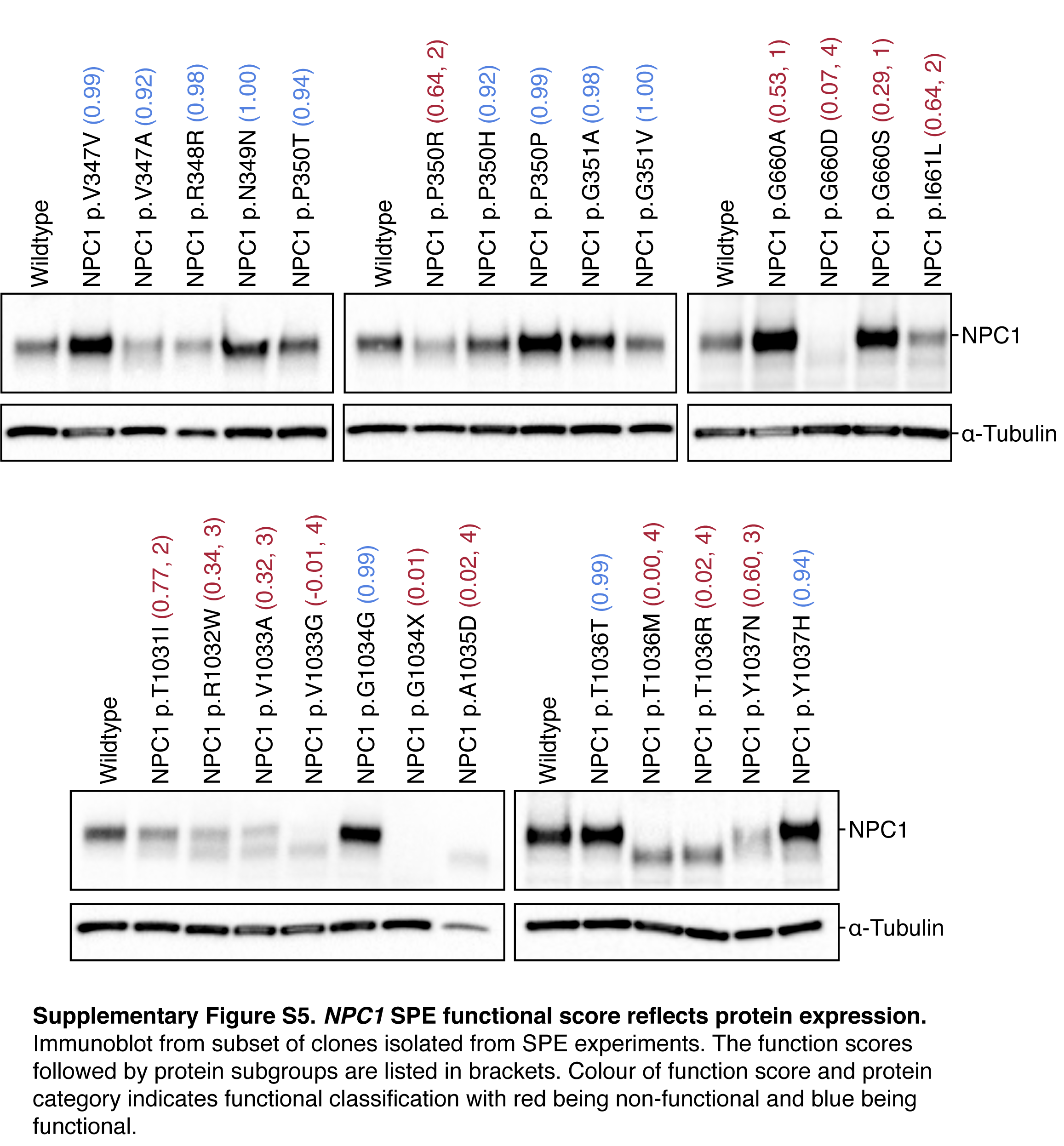

### Supplementary Figure S6

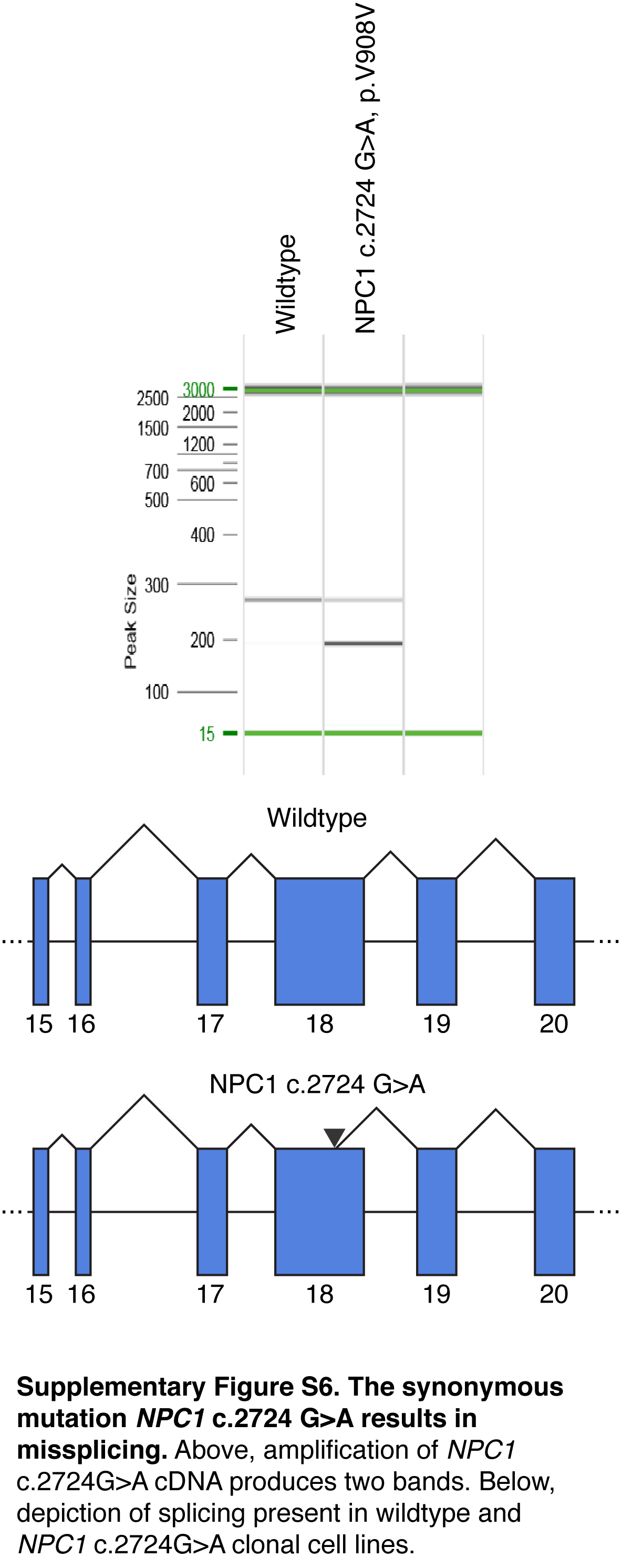
